# Visualizing the Structure of RNA-seq Expression Data using Grade of Membership Models

**DOI:** 10.1101/051631

**Authors:** Kushal K Dey, Chiaowen Joyce Hsiao, Matthew Stephens

**Affiliations:** Department of Statistics, University of Chicago, Chicago, Illinois 60637, USA; Department of Human Genetics, University of Chicago, Chicago, Illinois 60637, USA

## Abstract

Grade of membership models, also known as “admixture models”, “topic models” or “Latent Dirichlet Allocation”, are a generalization of cluster models that allow each sample to have membership in multiple clusters. These models are widely used in population genetics to model admixed individuals who have ancestry from multiple “populations”, and in natural language processing to model documents having words from multiple “topics”. Here we illustrate the potential for these models to cluster samples of RNA-seq gene expression data, measured on either bulk samples or single cells. We also provide methods to help interpret the clusters, by identifying genes that are distinctively expressed in each cluster. By applying these methods to several example RNA-seq applications we demonstrate their utility in identifying and summarizing structure and heterogeneity. Applied to data from the GTEx project on 53 human tissues, the approach highlights similarities among biologically-related tissues and identifies distinctively-expressed genes that recapitulate known biology. Applied to single-cell expression data from mouse preimplantation embryos, the approach highlights both discrete and continuous variation through early embryonic development stages, and highlights genes involved in a variety of relevant processes – from germ cell development, through compaction and morula formation, to the formation of inner cell mass and trophoblast at the blastocyst stage. The methods are implemented in the Bioconductor package CountClust.

**Author Summary:** Gene expression profile of a biological sample (either from single cells or pooled cells) results from a complex interplay of multiple related biological processes. Consequently, for example, distal tissue samples may share a similar gene expression profile through some common underlying biological processes. Our goal here is to illustrate that grade of membership (GoM) models – an approach widely used in population genetics to cluster admixed individuals who have ancestry from multiple populations – provide an attractive approach for clustering biological samples of RNA sequencing data. The GoM model allows each biological sample to have partial memberships in multiple biologically-distinct clusters, in contrast to traditional clustering methods that partition samples into distinct subgroups. We also provide methods for identifying genes that are distinctively expressed in each cluster to help biologically interpret the results. Applied to a dataset of 53 human tissues, the GoM approach highlights similarities among biologically-related tissues and identifies distinctively-expressed genes that recapitulate known biology. Applied to gene expression data of single cells from mouse preimplantation embryos, the approach highlights both discrete and continuous variation through early embryonic development stages, and genes involved in a variety of relevant processes. Our study highlights the potential of GoM models for elucidating biological structure in RNA-seq gene expression data.

## Introduction

Ever since large-scale gene expression measurements have been possible, clustering – of both genes and samples – has played a major role in their analysis [5–7]. For example, clustering of genes can identify genes that are working together or are co-regulated, and clustering of samples is useful for quality control as well as identifying biologically-distinct subgroups. A wide range of clustering methods have therefore been employed in this context, including distance-based hierarchical clustering, *k*-means clustering, and self-organizing maps (SOMs); see for example [8, 9] for reviews.

Here we focus on cluster analysis of samples, rather than clustering of genes (although our methods do highlight sets of genes that distinguish each cluster). Traditional clustering methods for this problem attempt to partition samples into distinct groups that show “similar” expression patterns. While partitioning samples in this way has intuitive appeal, it seems likely that the structure of a typical gene expression data set will be too complex to be fully captured by such a partitioning. Motivated by this, here we analyse expression data using grade of membership (GoM) models [10], which generalize clustering models to allow each sample to have partial membership in multiple clusters. That is, they allow that each sample has a proportion, or “grade” of membership in each cluster. Such models are widely used in population genetics to model admixture, where individuals can have ancestry from multiple populations [16], and in document clustering [41, 42] where each document can have membership in multiple topics. In these fields GoM models are often known as “admixture models”, and “topic models” or “Latent Dirichlet Allocation” [41]. GoM models have also recently been applied to detect mutation signatures in cancer samples [38].

Although we are not the first to apply GoM-like models to gene expression data, previous applications have been primarily motivated by a specific goal, “cell type deconvolution”, which involves using cell-type-specific expression profiles of marker genes to estimate the proportions of different cell types in a mixture [47, 49, 50]. Specifically, the GoM model we use here is analogous to – although different in detail from – blind deconvolution approaches [45, 46, 48] which estimate cell type proportions and cell type signatures jointly (see also [43, 44] for semi-supervised approaches). Our goal here is to demonstrate that GoM models can be useful much more broadly for understanding structure in RNA-seq data – not only to deconvolve mixtures of cell types. For example, in our analysis of human tissue samples from the GTEX project below, the GoM model usefully captures biological heterogeneity among samples even though the inferred grades of membership are unlikely to correspond precisely to proportions of specific cell types. And in our analyses of single-cell expression data the GoM model highlights interesting structure, even though interpreting the grades of membership as “proportions of cell types” is clearly inappropriate because each sample is a single cell! Here we are exploiting the GoM as a flexible extension of traditional cluster models, which can capture “continuous” variation among cells as well as the more “discrete” variation captured by cluster models. Indeed, the extent to which variation among cells can be described in terms of discrete clusters versus more continuous populations is a fundamental question that, when combined with appropriate single-cell RNA-seq data, the GoM models used here may ultimately help address.

## Methods Overview

We assume that the RNA-seq data on *N* samples has been summarized by a table of counts *C_N×G_ =* (*c_ng_*), where *c_ng_* is the number of reads from sample *n* mapped to gene *g* (or other unit, such as transcript or exon) [14]. The GoM model is a generalization of a cluster model, which allows that each sample has some proportion (“grade”) of membership, in each cluster. For RNA-seq data this corresponds to assuming that each sample *n* has some proportion of its reads, *q_nk_* coming from cluster *k*. In addition, each cluster *k* is characterized by a probability vector, *θ_k•_*, whose *g*th element represents the relative expression of gene *g* in cluster *k*. The GoM model is then

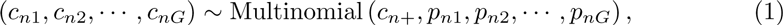

where

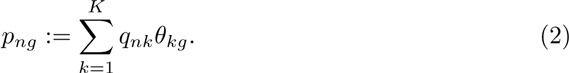

The number of clusters *K* is set by the analyst, and it can be helpful to explore multiple values of *K* (see Discussion).

To fit this model to RNA-seq data, we exploit the fact that this GoM model is commonly used for document clustering [41]. This is because, just as RNA-seq samples can be summarized by counts of reads mapping to each possible gene in the genome, document data can be summarized by counts of each possible word in a dictionary. Recognizing this allows existing methods and software for document clustering to be applied directly to RNA-seq data. Here we use the R package maptpx [15] to fit the GoM model.

Fitting the GoM model results in estimated membership proportions *q* for each sample, and estimated expression values *θ* for each cluster. We visualize the membership proportions for each sample using a “Structure plot” [17], which is named for its widespread use in visualizing the results of the *Structure* software [16] in population genetics. The Structure plot represents the estimated membership proportions of each sample as a stacked barchart, with bars of different colors representing different clusters. Consequently, samples that have similar membership proportions have similar amounts of each color. See Fig 1 for example.

**Fig 1.**
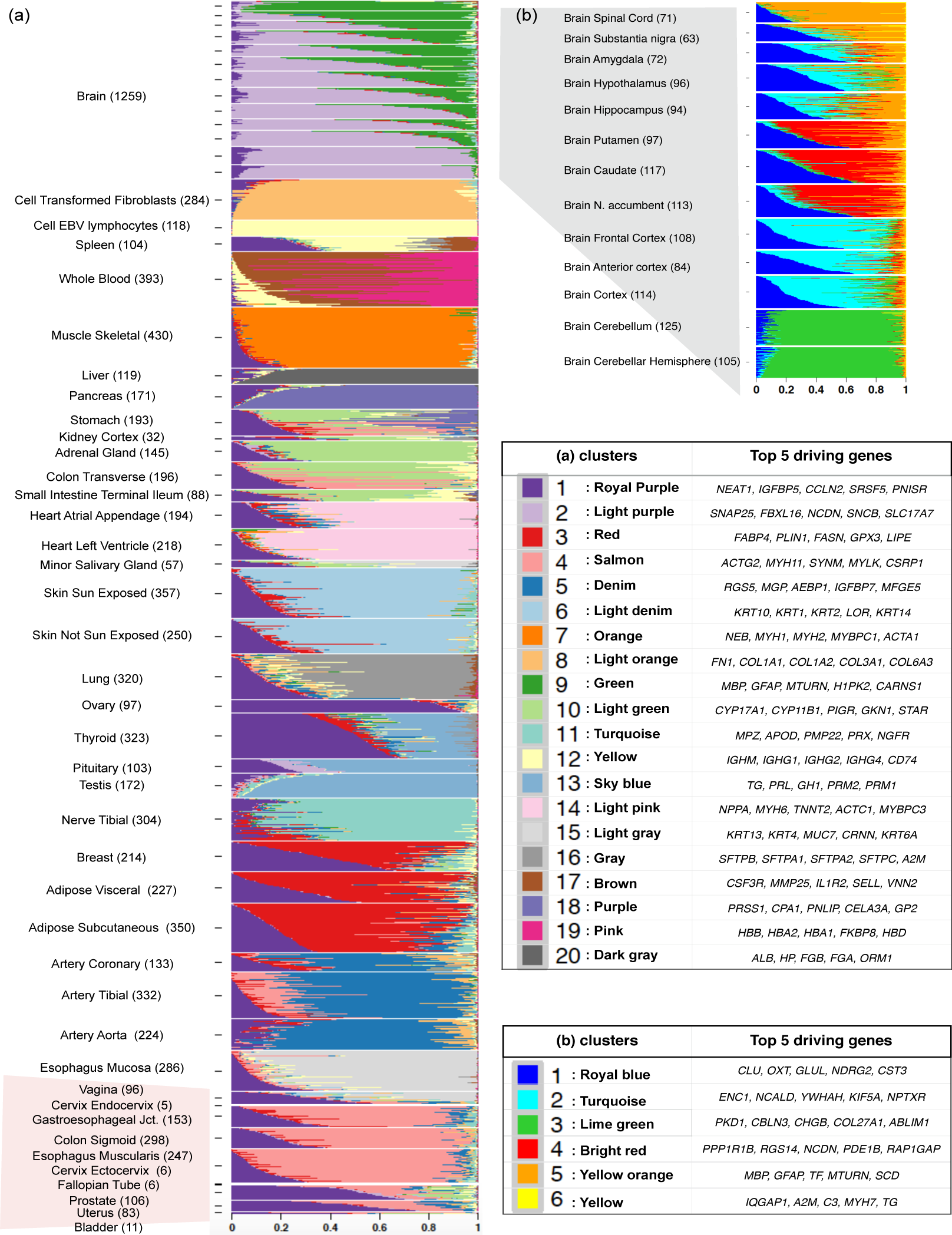
(a): Structure plot of estimated membership proportions for GoM model with *K* = 20 clusters fit to 8555 tissue samples from 53 tissues in GTEx data. Each horizontal bar shows the cluster membership proportions for a single sample, ordered so that samples from the same tissue are adjacent to one another. Within each tissue, the samples are sorted by the proportional representation of the underlying clusters.**(b)**: Structure plot of estimated membership proportions for *K* = 4 clusters fit to only the brain tissue samples. This analysis highlights finer-scale structure among the brain samples that is missed by the global analysis in (a).

To help biologically interpret the clusters inferred by the GoM model we also implemented methods to identify, for each cluster, which genes are most distinctively differentially expressed in that cluster; that is, which genes show the biggest difference in expression compared with the other most similar cluster (see Methods). Functions for fitting the GoM model, plotting the structure plots, and identifying the distinctive (“driving”) genes in each cluster, are included in our R package CountClust [53] available through Bioconductor [36].

## Results

### Bulk RNA-seq data of human tissue samples

We begin by illustrating the GoM model on bulk RNA expression measurements from the GTEx project (V6 dbGaP accession phs000424.v6.p1, release date: Oct 19, 2015, http://www.gtexportal.org/home/). These data consist of per-gene read counts from RNA-seq performed on 8,555 samples collected from 450 human donors across 53 tissues, lymphoblastoid cell lines, and transformed fibroblast cell-lines. We analyzed 16,069 genes that satisfied filters (e.g. exceeding certain minimum expression levels) that were used during eQTL analyses by the GTEx project (gene list available in http://stephenslab.github.io/count-clustering/project/utilities/gene_names_all_gtex.txt).

We fit the GoM model to these data, with number of clusters *K* = 5,10,15,20. For each *K* we ran the fitting algorithm three times and kept the result with the highest log-likelihood. As might be expected, increasing *K* highlights finer structure in the data, and for brevity we focus discussion on results for *K* = 20 (Fig 1(a)), with results for other *K* shown in S1 Fig. For comparison we also ran several other commonly-used methods for clustering and visualizing gene expression data: Principal Components Analysis (PCA), Multidimensional Scaling (MDS), *t*-Distributed Stochastic Neighbor Embedding (*t*-SNE) [25, 26], and hierarchical clustering (Fig 2).

**Fig 2.**
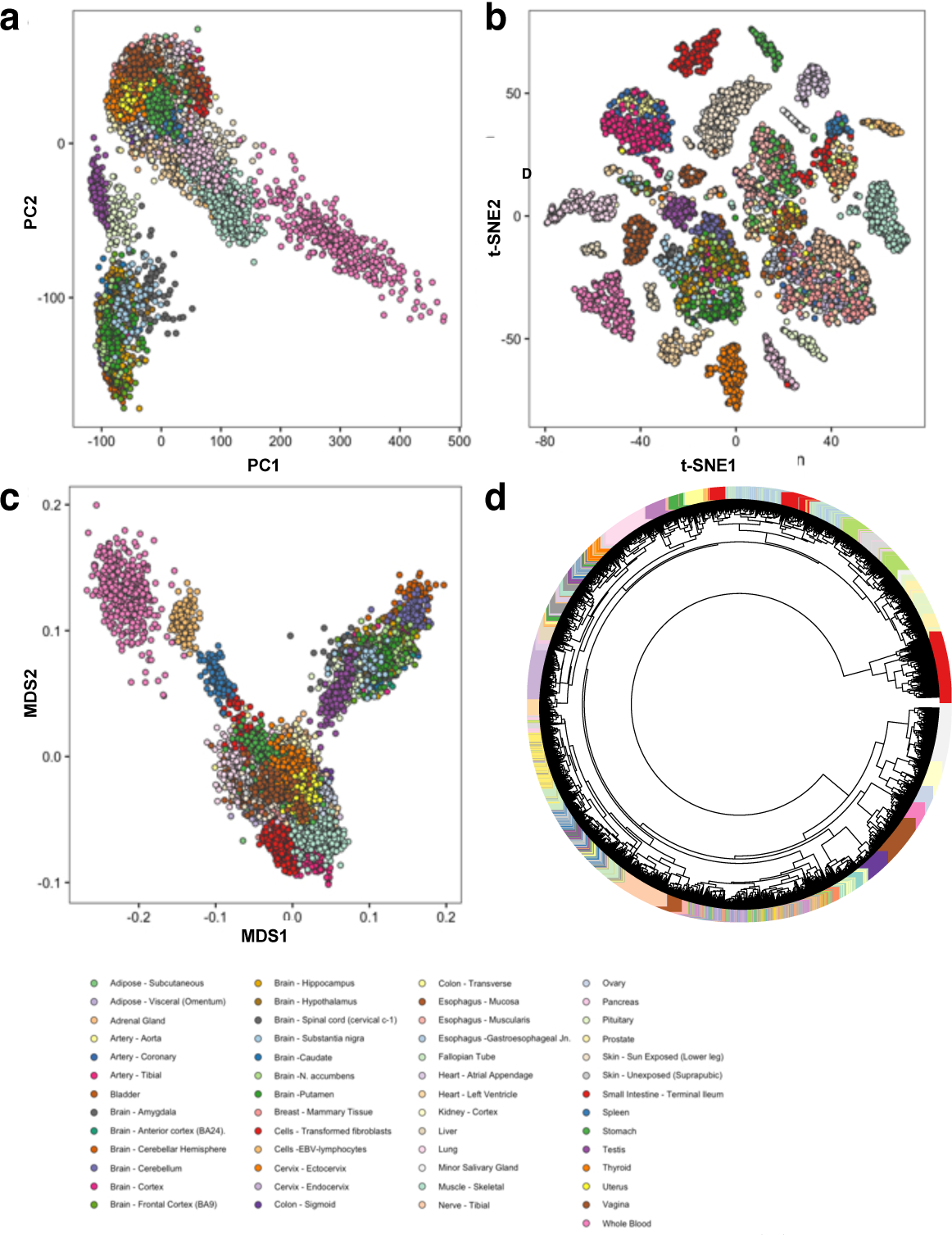
Visualization of the same GTEx data as in Figure 1 (a) across all tissues using standard and widely used approaches - Principal Component Analysis (PCA), Multi dimensional Scaling (MDS), t-SNE and hierarchical clustering. All the analysis are done on log CPM normalized expression data to remove library size effects. **(a)**: Plot of PC1 vs PC2 on the log CPM expression data, **(b)**: Plot of first two dimensions of the t-SNE plot, **(c)** Plot of first two dimensions of the Multi-Dimensional Scaling (MDS) plot. **(d)** Dendrogram for the hierarchical clustring of the GTEx tissue samples based on the log CPM expression data.

These data present a challenge to visualization and clustering tools, because of both the relatively large number of samples and the complex structure created by the inclusion of many different tissues. Indeed, neither PCA nor MDS provide satisfactory summaries of the structure in these data (Fig 2(a,b)): samples from quite different tissues are often super-imposed on one another in plots of PC1 vs PC2, and this issue is only partly alleviated by examining more PCs (Supplementary Figure S2 Fig). The hierarchical clustering provides perhaps better separation of tissues (Fig 2(d)), but producing a clear (static) visualization of the tree is difficult with this many samples. By comparison *t*-SNE (Fig 2(b)) and the GoM model (Fig 1(a)) both show a much clearer visual separation of samples by tissue, although they achieve this in very different ways. The *t*-SNE representation produces a two-dimensional plot with 20-25 visually-distinct clusters. In contrast, the GoM highlights similarity among samples by assigning them similar membership proportions, resulting in groups of similarly-colored bars in the structure plot. Some tissues are represented by essentially a single cluster/color (e.g. Pancreas, Liver), whereas other tissues are represented as a mixture of multiple clusters (e.g. Thyroid, Spleen). Furthermore, the GoM results highlight biological similarity among some tissues by assigning similar membership proportions to samples from those tissues. For example, samples from several different parts of the brain often have similar memberships, as do the arteries (aorta, tibial and coronary) and skin samples (sun-exposed and un-exposed).

Although it is not surprising that samples cluster by tissue, other results could have occurred. For example, samples could have clustered according to technical variables, such as sequencing batch [34] or sample collection center. While our results do not exclude the possibility that technical variables could have influenced these data, the *t*-SNE and GoM results clearly demonstrate that tissue of origin is the primary source of heterogeneity, and provide a useful initial assurance of data quality.

While in these data both the GoM model and *t*-SNE highlight the primary structure due to tissue of origin, the GoM results have at least two advantages over *t*-SNE. First, the GoM model provides an explicit, quantitative, estimate of the mean expression of each gene in each cluster, making it straightforward to assess which genes and processes drive differences among clusters; see Table 1 (and also S1 Table). Reassuringly, many results align with known biology. For example, the purple cluster (cluster 18), which distinguishes Pancreas from other tissues, is enriched for genes responsible for digestion and proteolysis, (e.g. *PRSS1*, *CPA1*, *PNLIP*). Similarly the yellow cluster (cluster 12), which primarily distinguishes Cell EBV Lymphocytes from other tissues, is enriched with genes responsible for immune responses (e.g. *IGHM*, *IGHG1*) and the pink cluster (cluster 19) which mainly appears in Whole Blood, is enriched with genes related hemoglobin complex and oxygen transport (e.g. *HBB*, *HBA1*, *HBA2*). Further, Keratin-related genes characterize the skin cluster (cluster 6, light denim), Myosin-related genes characterize the muscle skeletal cluster (cluster 7, orange), etc. These biological annotations are particularly helpful for understanding instances where a cluster appears in multiple tissues. For example, the top genes in the salmon cluster (cluster 4), which is common to the Gastroesophageal Junction, Esophagus Muscularis and Colon Sigmoid, are related to smooth muscle. And the top genes in the red cluster, highlighted above as common to Breast Mammary tissue, Adipose Subcutaneous and Adipose Visceral, are all related to adipocytes and/or fatty acid synthesis.

**Table 1.**
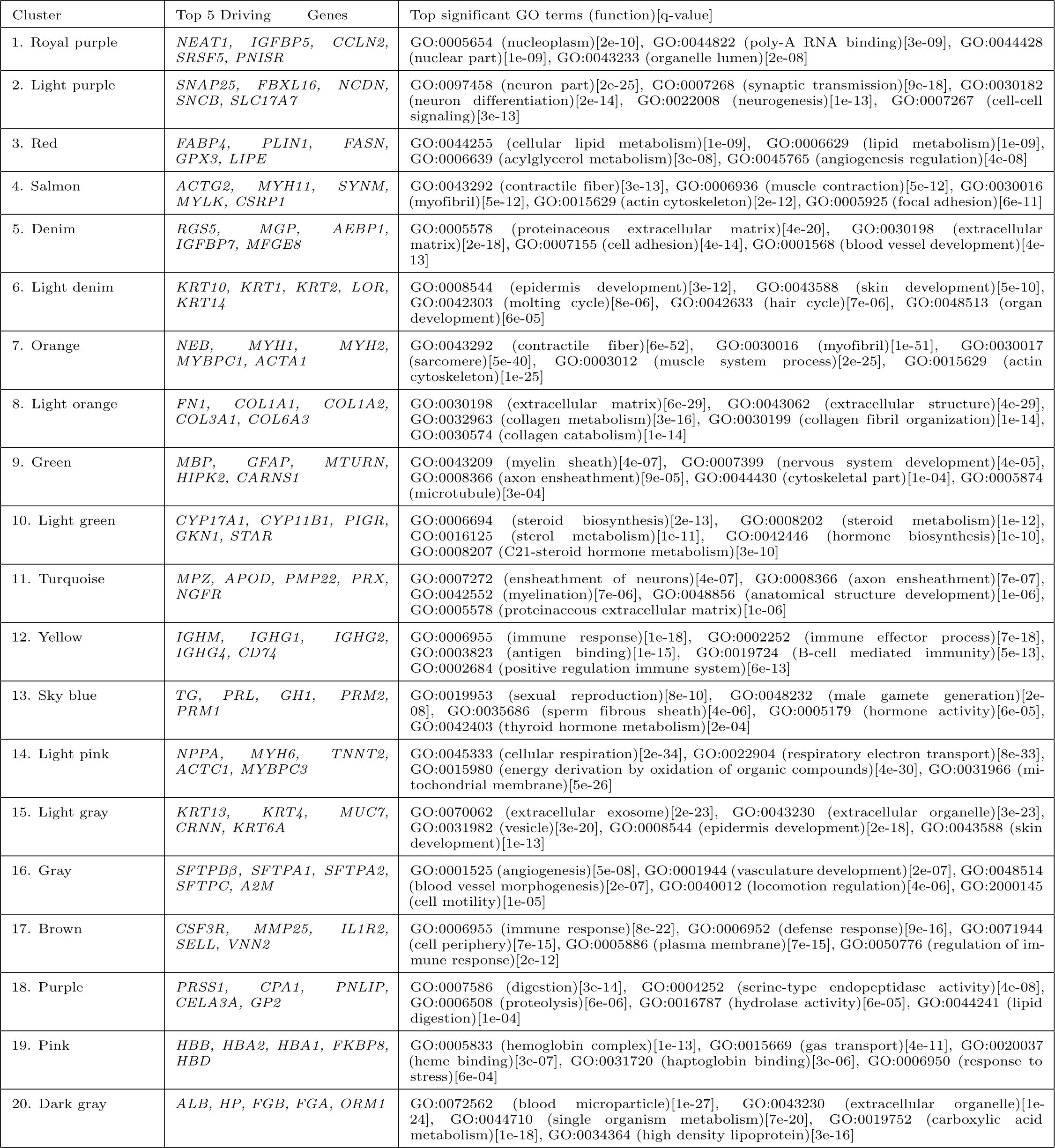
Cluster Annotations GTEx V6 data (with GO annotations).

A second advantage of the GoM model is that, because it allows partial membership in each cluster, it is better able to highlight partial similarities among distinct tissues. For example, in Figure 1(a) the sky blue cluster (cluster 13), appears in testis, pituitary, and thyroid, reflecting shared hormonal-related processes. At the same time, these tissues are distinguished from one another both by their degree of membership in this cluster (testis samples have consistently stronger membership; thyroid samples consistently weaker), and by membership in other clusters. For example, pituitary samples, but not testis or thyroid samples, have membership in the light purple cluster (cluster 2) which is driven by genes related to neurons and synapsis. In the *t*-SNE results these three tissues simply cluster separately into visually distinct groups, with no indication that their expression profiles have something in common (Fig 2(b)). Thus, although we find the *t*-SNE results visually attractive, this 2-dimensional projection contains less information than the Structure plot from the GoM (Fig 1(a)), which uses color to represent the samples in a 20-dimensional space.

In addition to these qualitative comparisons with other methods, we also used the GTEx data to quantitatively compare the accuracy of the GoM model with hierarchical clustering. Specifically, for each pair of tissues in the GTEx data we assessed whether or not each method correctly partitioned samples into the two tissue groups; see Methods. (Other methods do not provide an explicit clustering of the samples – only a visual representation – and so are not included in these comparisons.) The GoM model was more accurate in this test, succeeding in 88% of comparisons, compared with 79% for hierarchical clustering (Supplemental Figure S3 Fig (c) vs (a)).

### Sub-analysis of Brain tissues

Although the analysis of all tissues is useful for assessing global structure, it may miss finer-scale structure within tissues or among similar tissues. For example, here the GoM model applied to all tissues effectively allocated only three clusters to all brain tissues (clusters 1,2 and 9 in Fig 1(a)), and we suspected that additional substructure might be uncovered by analyzing the brain samples separately and using more clusters. Fig 1(b) shows the Structure plot for *K* = 6 on only the Brain samples. The results highlight much finer-scale structure compared with the global analysis. Brain Cerebellum and Cerebellar hemisphere are essentially assigned to a separate cluster (lime green), which is enriched with genes related to cell periphery and communication (e.g. *PKD1*, *CBLN3*) as well as genes expressed largely in neuronal cells and playing a role in neuron differentiation (e.g. *CHGB*). The spinal cord samples also show consistently strong membership in a single cluster (yellow-orange), the top defining gene for the cluster being *MBP* which is involved in myelination of nerves in the nervous system [51]. Another driving gene, *GFAP*, participates in system development by acting as a marker to distinguish astrocytes during development [4].

The remaining samples all show membership in multiple clusters. Samples from the putamen, caudate and nucleus accumbens show similar profiles, and are distinguished by strong membership in a cluster (cluster 4, bright red) whose top driving gene is *PPP1R1B*, a target for dopamine. And cortex samples are distinguished from others by stronger membership in a cluster (cluster 2, turquoise in Fig 1(b)) whose distinctive genes include *ENC1*, which interacts with actin and contributes to the organisation of the cytoskeleton during the specification of neural fate [3].

In comparison, applying PCA, MDS, hierarchical clustering and *t*-SNE to these brain samples reveals less of this finer-scale structure (Supplementary Figures S4 Fig). Both PCA and MDS effectively cluster the samples into two groups – those related to the cerebellum vs everything else. Hierarchical clustering also separates out the cerebellum-related tissues from the others, but again the format seems ill-suited to static visualization of more than one thousand samples. For reasons that we do not understand *t*-SNE performs poorly for these data: many samples are allocated to essentially identical locations, and so overplotting obscures them.

### Single-cell RNA-seq data

Recently RNA-sequencing has become viable for single cells [11], and this technology has the promise to revolutionize understanding of intra-cellular variation in expression, and regulation more generally [12]. Although it is traditional to describe and categorize cells in terms of distinct cell-types, the actual architecture of cell heterogeneity may be more complex, and in some cases perhaps better captured by the more “continuous” GoM model. In this section we illustrate the potential for the GoM model to be applied to single cell data.

To be applicable to single-cell RNA-seq data, methods must be able to deal with lower sequencing depth than in bulk RNA experiments: single-cell RNA-seq data typically involve substantially lower effective sequencing depth compared with bulk experiments, due to the relatively small number of molecules available to sequence in a single cell. Therefore, as a first step towards demonstrating its potential for single cell analysis, we checked robustness of the GoM model to sequencing depth. Specifically, we repeated the analyses above after thinning the GTEx data by a factor of 100 and 10,000 to mimic the lower sequencing depth of a typical single cell experiment. For the thinned GTEx data the Structure plot for *K* = 20 preserves most of the major features of the original analysis on unthinned data (Supplemental Figure S5 Fig). For the accuracy comparisons with hierarchical clustering, both methods suffer reduced accuracy in thinned data, but the GoM model remains superior (Supplemental Figure S6 Fig). For example, when thinning by a factor of 10,000, the success rate in separating pairs of tissues is 0.32 for the GoM model vs 0.10 for hierarchical clustering.

Having established its robustness to sequencing depth, we now illustrate the GoM model on two single cell RNA-seq datasets: data on mouse spleen from Jaitin *et al* [27] and data on mouse preimplantation embryos from Deng *et al* [28].

### Mouse Spleen data from Jaitin et al, 2014

Jaitin *et al* sequenced over 4,000 single cells from mouse spleen. Here we analyze 1,041 of these cells that were categorized as *CD*11*c*+ in the *sorting markers* column of their data (http://compgenomics.weizmann.ac.il/tanay/?page_id=519), and which had total number of reads mapping to non-ERCC genes greater than 600. Our hope was that applying the GoM model to these data would identify, and perhaps refine, the cluster structure evident in [27] (their Fig 2A and 2B). However, the GoM model yielded rather different results (Fig 3), where most cells were assigned to have membership in several clusters. Further, the cluster membership vectors showed systematic differences among amplification batches (which in these data is also strongly correlated with sequencing batch). For example, cells in batch 1 are characterized by strong membership in the orange cluster (cluster 5) while those in batch 4 are characterized by strong membership in both the blue and yellow clusters (2 and 6). Some adjacent batches show similar patterns - for example batches 28 and 29 have a similar visual “palette”, as do batches 32-45. And, more generally, these later batches are collectively more similar to one another than they are to the earlier batches (0-4).

**Fig 3.**
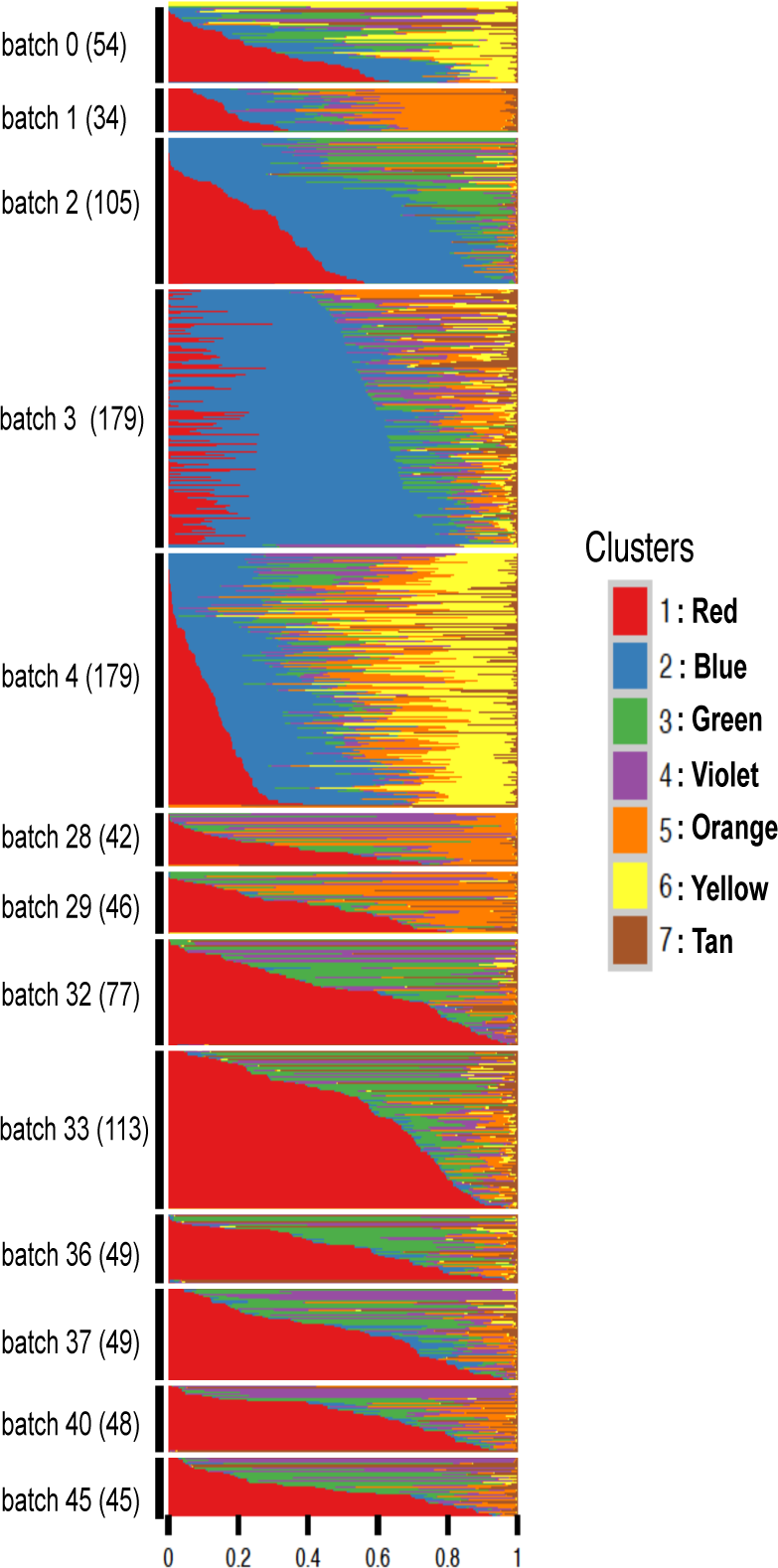
Structure plot of estimated membership proportions for GoM model with *K* = 7 clusters fit to 1,041 single cells from [27]. The samples (cells) are ordered so that samples from the same amplification batch are adjacent and within each batch, the samples are sorted by the proportional representation of the underlying clusters. In this analysis the samples do not appear to form clearly-defined clusters, with each sample being allocated membership in several “clusters”. Membership proportions are correlated with batch, and some groups of batches (e.g. 28-29; 32-45) show similar palettes. These results suggest that batch effects are likely influencing the inferred structure in these data.

The fact that batch effects are detectable in these data is not particularly surprising: there is a growing recognition of the importance of batch effects in high-throughput data generally [31, 32] and in single cell data specifically [33, 34]. And indeed, both clustering methods and the GoM model can be viewed as dimension reduction methods, and such methods can be helpful in controlling for batch effects [29, 30]. However, why these batch effects are not evident in Fig 2A and 2B of [27] is unclear.

### Mouse preimplantation embryo data from Deng et al, 2014

Deng *et al* collected single-cell expression data of mouse preimplantation embryos from the zygote to blastocyst stage [28], with cells from four different embryos sequenced at each stage. The original analysis [28] focuses on trends of allele-specific expression in early embryonic development. Here we use the GoM model to assess the primary structure in these data without regard to allele-specific effects (i.e. combining counts of the two alleles). Visual inspection of the Principal Components Analysis in [28] suggested perhaps 6-7 clusters, and we focus here on results with *K* = 6.

The results from the GoM model (Fig 4) clearly highlight changes in expression profiles that occur through early embryonic development stages, and enrichment analysis of the driving genes in each cluster (Table 3, S4 Table) indicate that many of these expression changes reflect important biological processes during embryonic preimplantation development.

**Fig 4.**
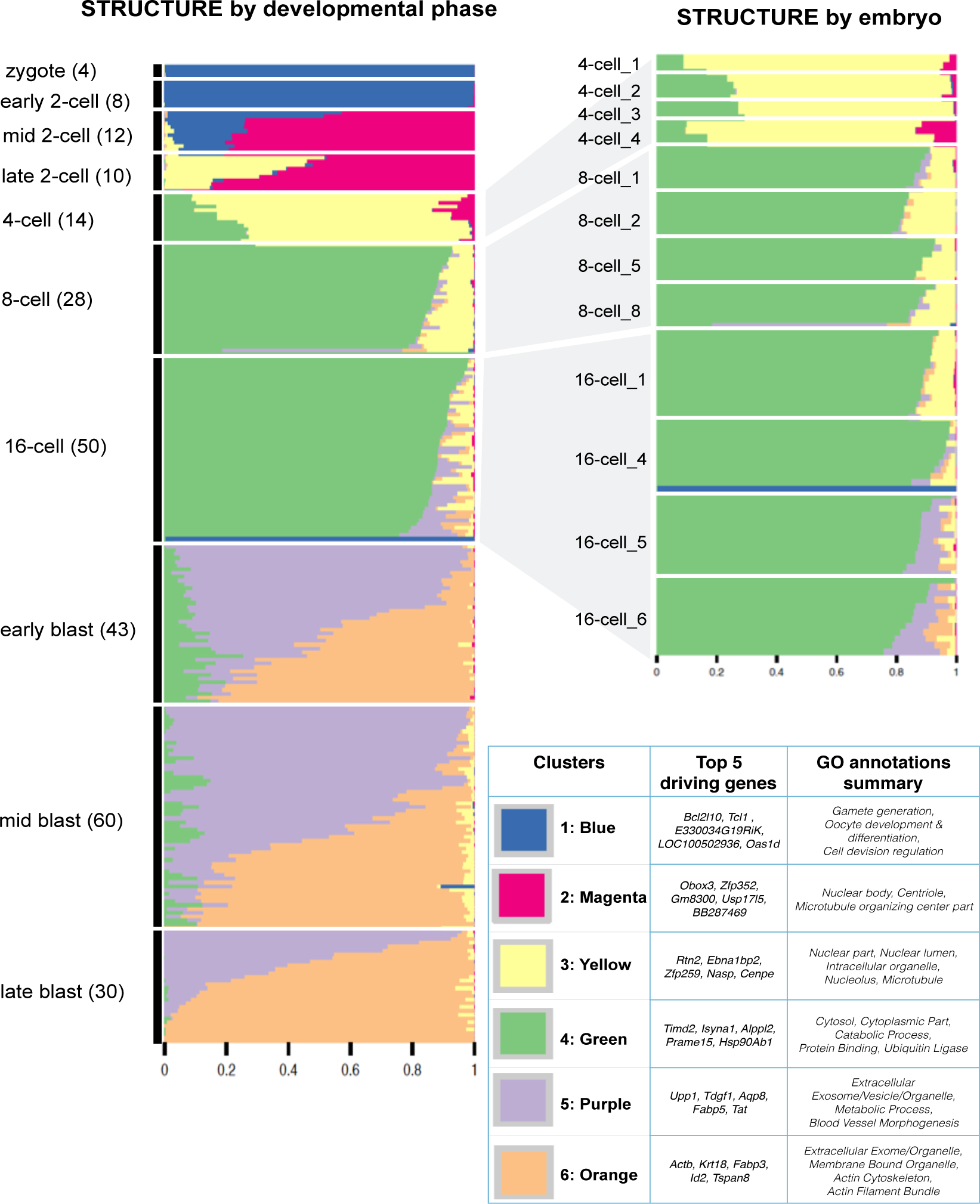
Structure plot of estimated membership proportions for GoM model with *K* = 6 clusters fit to 259 single cells from [28]. The cells are ordered by their preimplantation development phase (and within each phase, sorted by the proportional representation of the clusters). While the very earliest developmental phases (zygote and early 2-cell) are essentially assigned to a single cluster, others have membership in multiple clusters Each cluster is annotated by the genes that are most distinctively expressed in that cluster, and by the gene ontology categories for which these distinctive genes are most enriched (see Table 3 for more extensive annotation results). See text for discussion of biological processes driving these results.

In more detail: Initially, at the zygote and early 2-cell stages, the embryos are represented by a single cluster (blue in Fig 4) that is enriched with genes responsible for germ cell development (e.g., *Bcl2l10* [62], *Spin1* [63]). Moving through subsequent stages the grades of membership evolve to a mixture of blue and magenta clusters (mid 2-cell), a mixture of magenta and yellow clusters (late 2-cell) and a mixture of yellow and green (4-cell stage). The green cluster then becomes more prominent in the 8-cell and 16-cell stages, before dropping substantially in the early and mid-blastocyst stages. That is, we see a progression in the importance of different clusters through these stages, from the blue cluster, moving through magenta and yellow to green. Examining the genes distinguishing each cluster reveals that this progression (blue-magenta-yellow-green) reflects the changing relative importance of several fundamental biological processes. The magenta cluster is driven by genes responsible for the beginning of transcription of zygotic genes (e.g., *Zscan4c-f* show up in the list of top 100 driving genes: see https://stephenslab.github.io/count-clustering/project/src/deng_cluster_annotations.html), which takes place in the late 2-cell stage of early mouse embryonic development [65]. The yellow cluster is enriched for genes responsible for heterochromation *Smarcc1* [66] and chromosome stability *Cenpe* [67] (see S4 Table). And the green cluster is enriched for cytoskeletal genes (e.g., *Fbxo15*) and cytoplasm genes (e.g., *Tceb1*, *Hsp90ab1*), all of which are essential for compaction at the 8-cell stage and morula formation at the 16-cell stage.

Finally, during the blastocyst stages two new clusters (purple and orange in Fig 4) dominate. The orange cluster is enriched with genes involved in the formation of trophectoderm (TE) (e.g., *Tspan8*, *Krt8*, *Id2* [59]), while the purple cluster is enriched with genes responsible for the formation of inner cell mass (ICM) (e.g., *Pdgfra*,*Pyy* [61]).

For comparison, results for PCA, MDS, *t*-SNE and hierarchical clustering are shown in Supplemental Figure S7 Fig. All these methods show some clustering structure by pre-implantation stage; however only PCA and MDS seem to capture the developmental trajectory from zygote to blastocyst, exhibiting a “horse-shoe” pattern that is expected when similarities among samples approximately reflect an underlying latent ordering [39, 40]. And none of them provide any direct indication of the ICM vs TE structure in the blastocyst samples.

Although the GoM model results clearly highlight some of the key biological processes underlying embryonic preimplantation development, there are also some expected patterns that do not appear. Specifically, just prior to implantation the embryo consists of three different cell types, the trophectoderm (TE), the primitive endoderm (PE), and the epiblast (EPI) [60], with the PE and EPI being formed from the ICM. Thus one might expect the late blastocyst cells to show a clear division into three distinct groups, and for some of the earlier blastocyst cells to show partial membership in one of these groups as they begin to differentiate towards these cell types. Indeed, the GoM model seems well suited to capture this process in principle. However, this is not the result we obtained in practice. In particular, although the two clusters identified by the GoM model in the blastocyst stages appear to correspond roughly to the TE and ICM, even the late blastocyst cells tend to show a gradient of memberships in both these clusters, rather than a clear division into distinct groups. Our results contrast with those from the single-cell mouse preimplantation data of [59], measured by qPCR, where the late blastocyst cells showed a clear visual division into three groups using PCA (their Figure 1).

To better understand the differences between our results for RNA-seq data from [28] and the qPCR results from [59] we applied the GoM model with *K* = 3 to a small subset of the RNA-seq data: the blastocyst cell data at the 48 genes assayed by [59]. These genes were specifically chosen by them to help elucidate cell-fate decisions during early development of the mouse embryo. Still, the GoM model results (Supplemental Figure S8 Fig) do not support a clear division of these data into three distinct groups (and neither do PCA or *t*-SNE; Supplemental Figure S9 Fig). Rather, the GoM model highlights one cluster (Green in figure), whose membership proportions essentially reflect expression at the *Actb* gene, and two other clusters (Orange and Purple in figure) whose driving genes correspond to genes identified in [59] as being distinctive to TE and EPI cell types respectively. The *Actb* gene is a “housekeeping gene”, used by [59] to normalize their qPCR data, and its prominence in the GoM results likely reflects its very high expression levels relative to other genes. However, excluding *Actb* from the analysis still does not lead to a clear separation into three groups (Supplemental Figure S8 Fig). Thus, although there are clear commonalities in the structure of these RNA-seq and qPCR data sets, the structure of the single-cell RNA-seq data from [28] is fundamentally more complex (or, perhaps, muddied), and consequently more difficult to interpret.

In addition to trends across development stages, the GoM results also highlight some embryo-level effects in the early stages (Fig 4). Specifically, cells from the same embryo sometimes show greater similarity than cells from different embryos. For example, while all cells from the 16-cell stage have high memberships in the green cluster, cells from two of the embryos at this stage have memberships in both the purple and yellow clusters, while the other two embryos have memberships only in the yellow cluster.

The GoM results also highlight a few single cells as outliers. For example, a cell from a 16-cell embryo is represented by the blue cluster - a cluster that represents cells at the zygote and early 2-cell stage. Also, a cell from an 8-stage embryo has strong membership in the purple cluster - a cluster that represents cells from the blastocyst stage. This illustrates the potential for the GoM model to help in quality control: it would seem prudent to consider excluding these outlier cells from subsequent analyses of these data.

## Discussion

Our goal here is to highlight the potential for GoM models to elucidate structure in RNA-seq data from both single cell sequencing and bulk sequencing of pooled cells. We also provide tools to identify which genes are most distinctively expressed in each cluster, to aid interpretation of results. As our applications illustrate, the results can provide a richer summary of the structure in RNA-seq data than existing widely-used visualization methods such as PCA and hierarchical clustering. While it could be argued that the GoM model results sometimes raise more questions than they answer, this is exactly the point of an exploratory analysis tool: to highlight issues for investigation, identify anomalies, and generate hypotheses for future testing.

Our results from different methods also highlight another important point: different methods have different strengths and weaknesses, and can compliment one another as well as competing. For example, *t*-SNE seems to provide a much clearer indication of the cluster structure in the full GTEx data than does PCA, but does a poorer job of capturing the ordering of the developmental samples from mouse pre-implantation embryos. While we believe the GoM model often provides a richer summary of the sample structure, we would expect to use it in addition to *t*-SNE and PCA when performing exploratory analyses. (Indeed the methods can be used in combination: both PCA and *t*-SNE can be used to visualize the results of the GoM model, as an alternative or complement to the Structure plot.)

A key feature of the GoM model is that it allows that each sample has a proportion of membership in each cluster, rather than a discrete cluster structure. Consequently it can provide insights into how well a particular dataset really fits a “discrete cluster” model. For example, consider the results for the data from Jaitin *et al* [27] and Deng *et al* [28]: in both cases most samples are assigned to multiple clusters, although the results are closer to “discrete” for the latter than the former. The GoM model is also better able to represent the situation where there is not really a single clustering of the samples, but where samples may cluster differently at different genes. For example, in the GTEx data, the stomach samples share memberships in common with both the pancreas (purple) and the adrenal gland (light green). This pattern can be seen in the Structure plot (Fig 1) but not from other methods like PCA, *t*-SNE or hierarchical clustering (Fig 2).

Fitting GoM models can be computationally-intensive for large data sets. For the datasets we considered here the computation time ranged from 12 minutes for the data from [28] (*n* = 259; *K* = 6), through 33 minutes for the data from [27] (*n* = 1,041; *K* = 7) to 3,370 minutes for the GTEx data (*n* = 08,555; *K* = 20). Computation time can be reduced by fitting the model to only the most highly expressed genes, and we often use this strategy to get quick initial results for a dataset. Because these methods are widely used for clustering very large document datasets there is considerable ongoing interest in computational speed-ups for very large datasets, with “on-line” (sequential) approaches capable of dealing with millions of documents [56] that could be useful in the future for very large RNA-seq datasets.

A thorny issue that arises when fitting clustering models is how to select the number of clusters, *K*. Like many software packages for fitting these models, the maptpx package implements a measure of model fit that provides one useful guide. However, it is worth remembering that in practice there is unlikely to be a “true” value of *K*, and results from different values of *K* may complement one another rather than merely competing with one another. For example, seeing how the fitted model evolves as *K* increases is one way to capture some notion of hierarchy in the clusters identified [17]. More generally it is often fruitful to analyse data in multiple ways using the same tool: for example our GTEx analyses illustrate how analysis of subsets of the data (in this case the brain samples) can complement analyses of the entire data. Finally, as a practical matter, we note that Structure plots can be difficult to read for large *K* (e.g. *K* = 30) because of the difficulties of choosing a palette with *K* distinguishable colors.

The version of the GoM model fitted here is relatively simple, and could certainly be embellished. For example, the model allows the expression of each gene in each cluster to be a free parameter, whereas we might expect expression of most genes to be “similar” across clusters. This is analogous to the idea in population genetics applications that allele frequencies in different populations may be similar to one another [20], or in document clustering applications that most words may not differ appreciably in frequency in different topics. In population genetics applications incorporating this idea into the model, by using a correlated prior distribution on these frequencies, can help improve identification of subtle structure [20] and we would expect the same to happen here for RNA-seq data.

Finally, GoM models can be viewed as one of a larger class of “matrix factorization” approaches to understanding structure in data, which also includes PCA, non-negative matrix factorization (NMF), and sparse factor analysis (SFA); see [21]. This observation raises the question of whether methods like SFA might be useful for the kinds of analyses we performed here. (NMF is so closely related to the GoM model that we do not discuss it further; indeed, the GoM model is a type of NMF, because both grades of membership and expression levels within each cluster are required to be non-negative.) Informally, SFA can be thought of as a generalization of the GoM model that allows samples to have *negative* memberships in some “clusters” (actually, “factors”). This additional flexibility should allow SFA to capture certain patterns more easily than the GoM model. For example, a small subset of genes that are over-expressed in some samples and under-expressed in other samples could be captured by a single sparse factor, with positive loadings in the over-expressed samples and negative loadings in the other samples. However, this additional flexibility also comes at a cost of additional complexity in visualizing the results. For example, Supplementary Figures S10 Fig, S11 Fig, S12 Fig show results of SFA (the version from [21]) for the GTEx data and the mouse preimplantation data: in our opinion, these do not have the simplicity and immediate visual appeal of the GoM model results. Also, applying SFA to RNA-seq data requires several decisions to be made that can greatly impact the results: what transformation of the data to use; what method to induce sparsity (there are many; e.g.[21–24]); whether to induce sparsity in loadings, factors, or both; etc. Nonetheless, we certainly view SFA as complementing the GoM model as a promising tool for investigating the structure of RNA-seq data, and as a promising area for further work.

## Methods and Materials

### Model Fitting

We use the maptpx R package [15] to fit the GoM model (1,2), which is also known as “Latent Dirichlet Allocation” (LDA). The maptpx package fits this model using an EM algorithm to perform Maximum a posteriori (MAP) estimation of the parameters *q* and *θ*. See [15] for details.

### Visualizing Results

In addition to the Structure plot, we have also found it useful to visualize results using t-distributed Stochastic Neighbor Embedding (t-SNE), which is a method for visualizing high dimensional datasets by placing them in a two dimensional space, attempting to preserve the relative distance between nearby samples [25, 26]. Compared with the Structure plot our t-SNE plots contain less information, but can better emphasize clustering of samples that have similar membership proportions in many clusters. Specifically, t-SNE tends to place samples with similar membership proportions together in the two-dimensional plot, forming visual “clusters” that can be identified by eye (e.g.http://stephenslab.github.io/count-clustering/project/src/tissues_tSNE_2.html). This may be particularly helpful in settings where no external information is available to aid in making an informative Structure plot.

### Cluster annotation

To help biologically interpret the clusters, we developed a method to identify which genes are most distinctively differentially expressed in each cluster. (This is analogous to identifying “ancestry informative markers” in population genetics applications [18].) Specifically, for each cluster *k* we measure the distinctiveness of gene *g* with respect to any other cluster *l* using

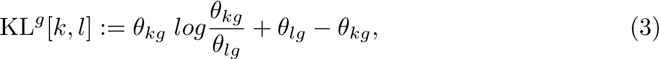

which is the Kullback–Leibler divergence of the Poisson distribution with parameter *θ_kg_* to the Poisson distribution with parameter *θ_lg_*. For each cluster *k*, we then define the distinctiveness of gene *g* as

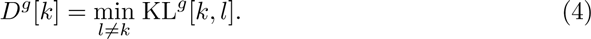

The higher *D^g^*[*k*], the larger the role of gene *g* in distinguishing cluster *k* from all other clusters. Thus, for each cluster *k* we identify the genes with highest *D^g^*[*k*] as the genes driving the cluster *k*. We annotate the biological functions of these individual genes using the mygene R Bioconductor package [35].

For each cluster *k*, we filter out a number of genes (top 100 for the Deng *et al* data [28] and GTEx V6 data [13]) with highest *D^g^*[*k*] value and perform a gene set over-representation analysis of these genes against all the other genes in the data representing the background. To do this, we used ConsensusPathDB database (http://cpdb.molgen.mpg.de/) [57] [58]. See Table 1-2 and Table 3 for the top significant gene ontologies driving each cluster in the GTEx V6 data and the Deng *et al*data respectively.

**Table 2.** Cluster Annotations for GTEx V6 Brain data.

| Cluster | Top 5 Driving Genes | Top significant GO terms |
| --- | --- | --- |
| 1. Royal blue | *CLU, OXT, GLUL, NDRG2, CST3* | GO:0043230 (extracellular organelle)[5e-11], GO:1903561 (extracellular vesicle)[6e-11], GO:0070062 (extracellular exosome)[2e-09], GO:0006950 (response to stress)[3e-10], GO:0031988 (membrane bound vesicle)[1e-10] |
| 2. Turquoise | *ENC1, NCALD, YWHAH, KIF5A, NPTXR* | GO:0097458 (neuron part)[3e-11], GO:0008092 (cytoskeletal protein binding)[7e-11], GO:0031175 (neuron projection development)[7e-09], GO:0030182 (neuron differentiation)[4e-08], GO:0007268 (synaptic transmission)[1e-08] |
| 3. Lime green | *PKD1, CBLN3, CHGB, COL27A1, ABLIM1* | GO:0005089 (Rho guanyl-nucleotide exchange factor activity)[1e-03], GO:0016604 (nuclear body)[0.002], GO:0022008 (neurogenesis)[0.02], GO:0035239 (tube morphogenesis)[0.08], GO:0007269 (neurotransmitter secretion)[0.10] |
| 4. Red | *PPP1R1B, RGS14, NCDN, PDE1B, RAP1GAP* | GO:0065009 (regulation of molecular function)[2e-06], GO:0036477 (somatodendritic compartment)[6e-05], GO:0007268 (synaptic transmission)[1e-03], GO:0023051 (signaling regulation)[2e-03], GO:0010646 (cell communication regulation)[1e-03] |
| 5. Yellow orange | *MBP, GFAP, TF, MTURN, SCD* | GO:0043209 (myelin sheath)[2e-09], GO:0007399 (nervous system development)[1e-04], GO:0005737 (cytoplasm)[1e-04], GO:0048471 (perinuclear region of cytoplasm)[5e-04], GO:0007272 (ensheathment of neurons)[1e-02] |
| 6. Yellow | *IQGAP1, A2M, C3, MYH7, TG* | GO:0072562 (blood microparticle)[1e-10], GO:0044449 (contractile fiber part)[1e-10], GO:0043230 (extracellular organelle)[7e-10], GO:0030017 (sarcomere)[1e-08], GO:0072376 (protein activation cascade)[1e-08] |

**Table 3.**
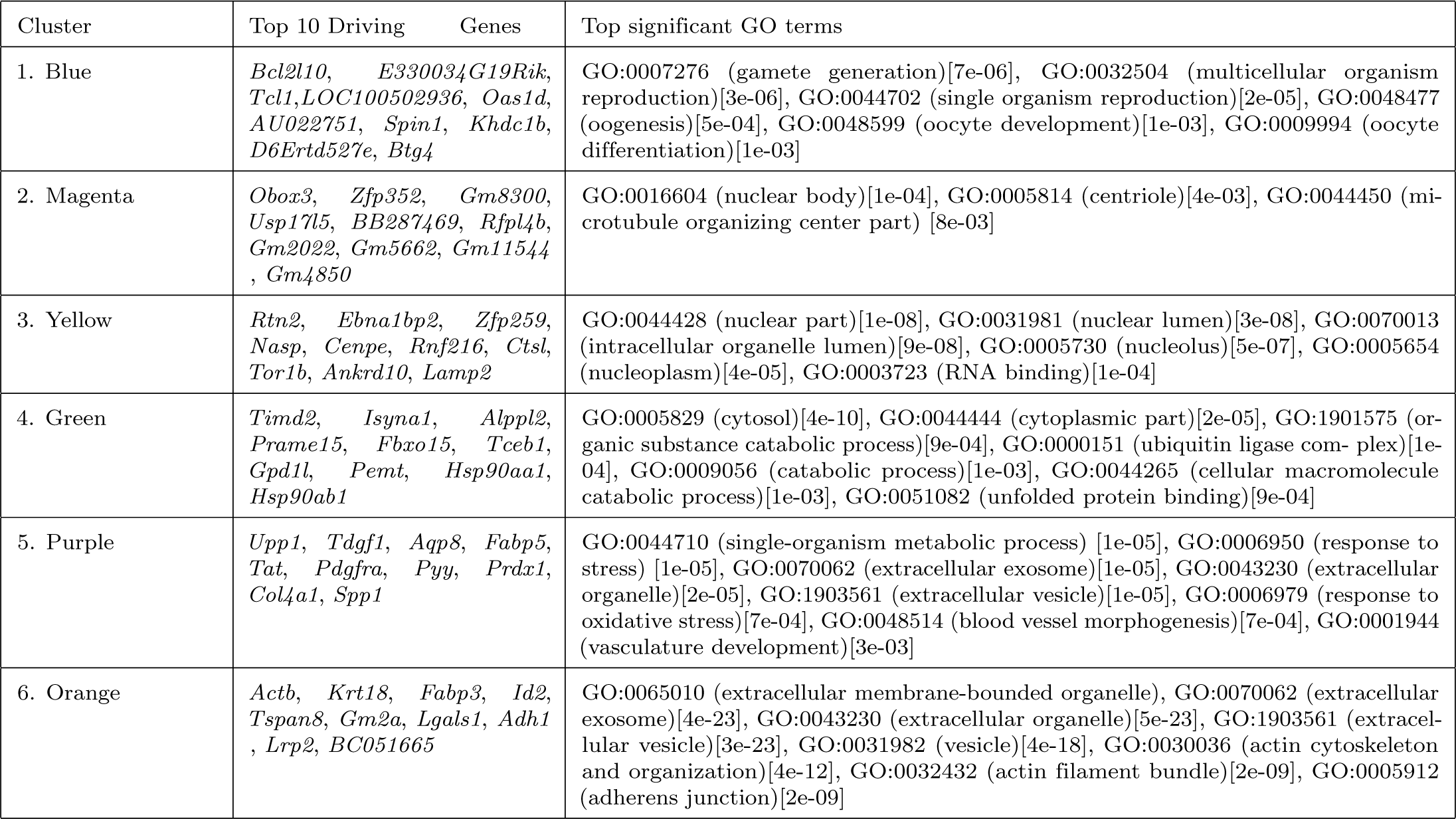
Cluster Annotations for Deng et al (2014) data.

### Comparison with hierarchical clustering

We compared the GoM model with a distance-based hierarchical clustering algorithm by applying both methods to samples from pairs of tissues from the GTEx project, and assessed their accuracy in separating samples according to tissue. For each pair of tissues we randomly selected 50 samples from the pool of all samples coming from these tissues. For the hierarchical clustering approach we cut the dendrogram at *K* = 2, and checked whether or not this cut partitions the samples into the two tissue groups. (We applied hierarchical clustering using Euclidean distance, with both complete and average linkage; results were similar and so we showed results only for complete linkage.)

For the GoM model we analysed the data with *K* = 2, and sorted the samples by their membership in cluster 1. We then partitioned the samples at the point of the steepest fall in this membership, and again we checked whether this cut partitions the samples into the two tissue groups. Supplemental Figure S3 Fig shows, for each pair of tissues, whether each method successfully partitioned the samples into the two tissue groups.

### Thinning

We used “thinning” to simulate lower-coverage data from the original higher-coverage data.. Specifically, if *c_ng_* is the counts of number of reads mapping to gene *g* for sample *n* for the original data, we simulated thinned counts *t_ng_* using

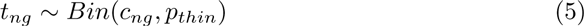

where *p_thin_* is a specified thinning parameter.

### Code Availability

Our methods are implemented in an R package CountClust, available as part of the Bioconductor project at https://www.bioconductor.org/packages/3.3/bioc/html/CountClust.html. The development version of the package is also available at https://github.com/kkdey/CountClust.

Code for reproducing results reported here is available at http://stephenslab.github.io/count-clustering/.

## Acknowledgments

We thank Matt Taddy, Amos Tanay and Effi Kenigsberg for helpful discussions. We thank Po-Yuan Tung, John Blischak and Jonathan Pritchard for helpful comments on the draft manuscript.

## Supporting Information

**S1 Fig. Structure plot of GTEx V6 tissue samples for (a) *K* = 5, (b) *K* = **10, (c)** *K* = 15, (d) *K* = 20.**Some tissues form a separate cluster from the other tissues from *K* = 5 onwards (for example: Whole Blood, Skin), whereas some tissue only form a distinctive subgroup at *K* = 20 (for example: Arteries).

**S2 Fig. Top five principal components (PC) for GTEx V6 tissue samples.** Scatter plot representation of the top five PCs of the GTEx tissue samples. Data was transformed to log2 counts per million (CPM).

**S3 Fig. Comparison between GoM model and hierarchical clustering under different scenarios of data transformation.** We used GTEx V6 data for model performance comparisons. Specifically, for every pair of the 53 tissues, we assessed the ability of the methods to separate samples according to their tissue of origin. The subplots of heatmaps show the results of evaluation under different scenarios. Filled squares in the heatmap indicate successful separation of the samples in corresponding tissue pair comparison. (a) Hierarchical clustering on log2 counts per million (CPM) transformed data using Euclidean distance. (b) Hierarchical clustering on the standardized log2-CPM transformed data (transformed values for each gene was mean and scale transformed) using the Euclidean distance. (c) GoM model of *K* = 2 applied to counts. (d) Hierarchical clustering on counts data with the assumption that, for each gene the sample read count *c_ng_* has a variance *c*¯*_g_ +* 1 that is constant across samples. And, the the gene-specific variance *c*¯*_g_ +* 1 was used to scale the distance matrix for clustering. (e) Hierarchical clustering applied to adjusted count data. Each gene has a mean expression value of 0 and variance of 1. Taken together, these results suggest that regardless of the different data transformation scenarios, the GoM model with *K =* 2 is able to separate samples of different tissue of origin, better than hierarchical cluster methods.

**S4 Fig. GTEx brain PCA, t-SNE and MDS.**

**S5 Fig. Structure plot of GTEx V6 tissue samples for *K* = 20 in two runs under different thinning parameter settings.** (a) *p_thin_ =* 0.01 and (B)*p_thin_ =* 0.0001. The structure in these two plots closely resemble the pattern observed in Fig 1(a), though there are a few differences from the unthinned version.

**S6 Fig. A comparison of accuracy of hierarchical clustering vs GoM on thinned GTEx data, with thinning parameters of** *p_thin_ =* 0.01 **and** *p_thin_ =* 0.001. For each pair of tissue samples from the GTEx V6 data we assessed whether or not each clustering method (with *K* = 2 clusters) separated the samples according to their tissue of origin, with successful separation indicated by a filled square. Thinning deteriorates accuracy compared with the unthinned data (Fig 2), but even then the model-based method remains more successful than the hierarchical clustering in separating the samples by tissue or origin.

**S7 Fig. Deng et al (2014) PCA, tSNE, MDS and dendrogram plots for hierarchical clustering.**

**S8 Fig. Additional GoM analysis of Deng et al (2014) data including blastocyst samples and 48 blastocyst marker genes.** We considered 48 blastocyst marker genes (as chosen by Guo et al., 2010) and fitted GoM model with *K*= 3 to 133 blastocyst samples. In the Structure plot, blastocyst samples are arranged in order of estimated membership proportion in the Green cluster. The panel located above the Structure plot shows the corresponding pre-implantation stage from which blastocyst samples were collected. The heatmap located below the Structure plot represents expression levels of the 48 blastocyst marker genes (log2 CPM), and the corresponding dendrogram shows results of hierarchical clustering (complete linkage). The table on the right of the expression heatmap displays gene information, showing, from left to right, 1) whether or not the gene is a transcription factor, 2) the driving GoM cluster if the gene was among the top five driving genes, and 3) the featured cell type (TE: trophecoderm, EPI: epiblast, PE: primitive endoderm) that was found in Guo et al., 2010.

**S9 Fig. Visualization of PCA and t-SNE results of mouse pre-implantation embryos data from Deng et al (2014) using 48 blastocyst marker genes.**

**S10 Fig. Sparse Factor Analysis loadings visualization of GTEx V6 tissue samples.** The colors represent the 20 different factors. The factor loadings are presented in a stacked bar for each sample. We performed SFA under the scenarios of when the loadings are sparse (left panel) and when the factors are sparse (right panel).

**S11 Fig. Sparse Factor Analysis loadings visualization of GTEx brain tissue samples.** The colors represent the 6 different factors. The factor loadings are presented in a stacked bar for each sample. We performed SFA under the scenarios of when the loadings are sparse (left panel) and when the factors are sparse (right panel).

**S12 Fig. Sparse Factor Analysis loadings visualization of mouse pre-implantation embryos from Deng et al., (2014). The colors represent the 6 different factors. The factor loadings are presented in a stacked bar for each sample. We performed SFA under the scenarios of when the loadings are sparse (left panel) and when the factors are sparse (right panel).**

**S1 Table. Cluster Annotations of GTEx V6 data with top driving gene summaries.**

**S2 Table. Cluster Annotations of GTEx V6 Brain data with top driving gene summaries.**

**S3 Table. Cluster Annotations of Deng data with top driving genes.**

**S4 Table. Cluster Annotation of Deng data analysis using 48 genes with top driving gene summaries.**

## Supplementary figures

**S1 Fig.**
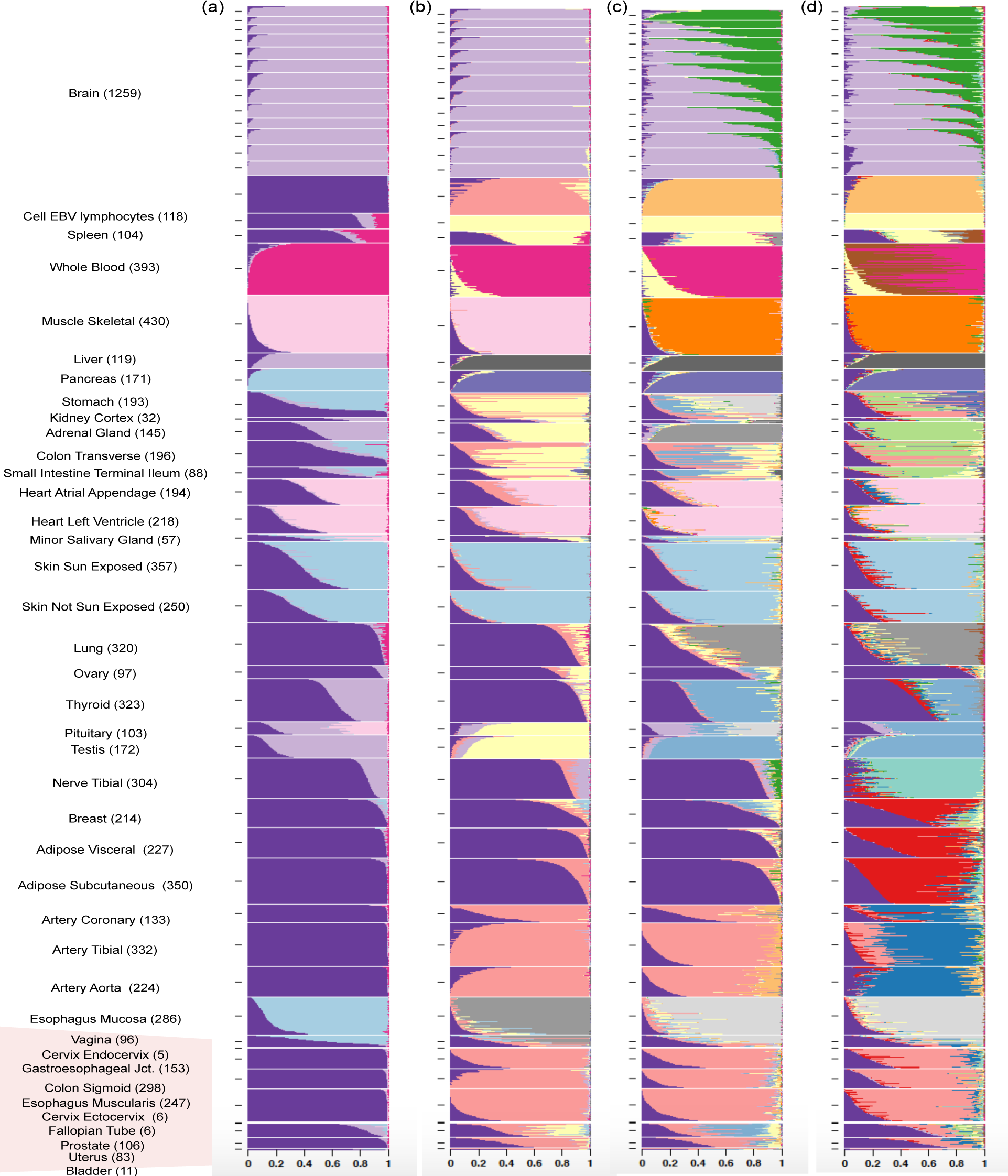
Structure plot of GTEx V6 tissue samples for (A) *K* = 5, (B) *K* = 10, (C) *K* = 15,(D) *K* = 20. Some tissues form a separate cluster from the other tissues from *K* = 5 onwards (for example: Whole Blood, Skin), whereas some tissue only form a distinctive subgroup at *K* = 20 (for example: Arteries).

**S2 Fig.**
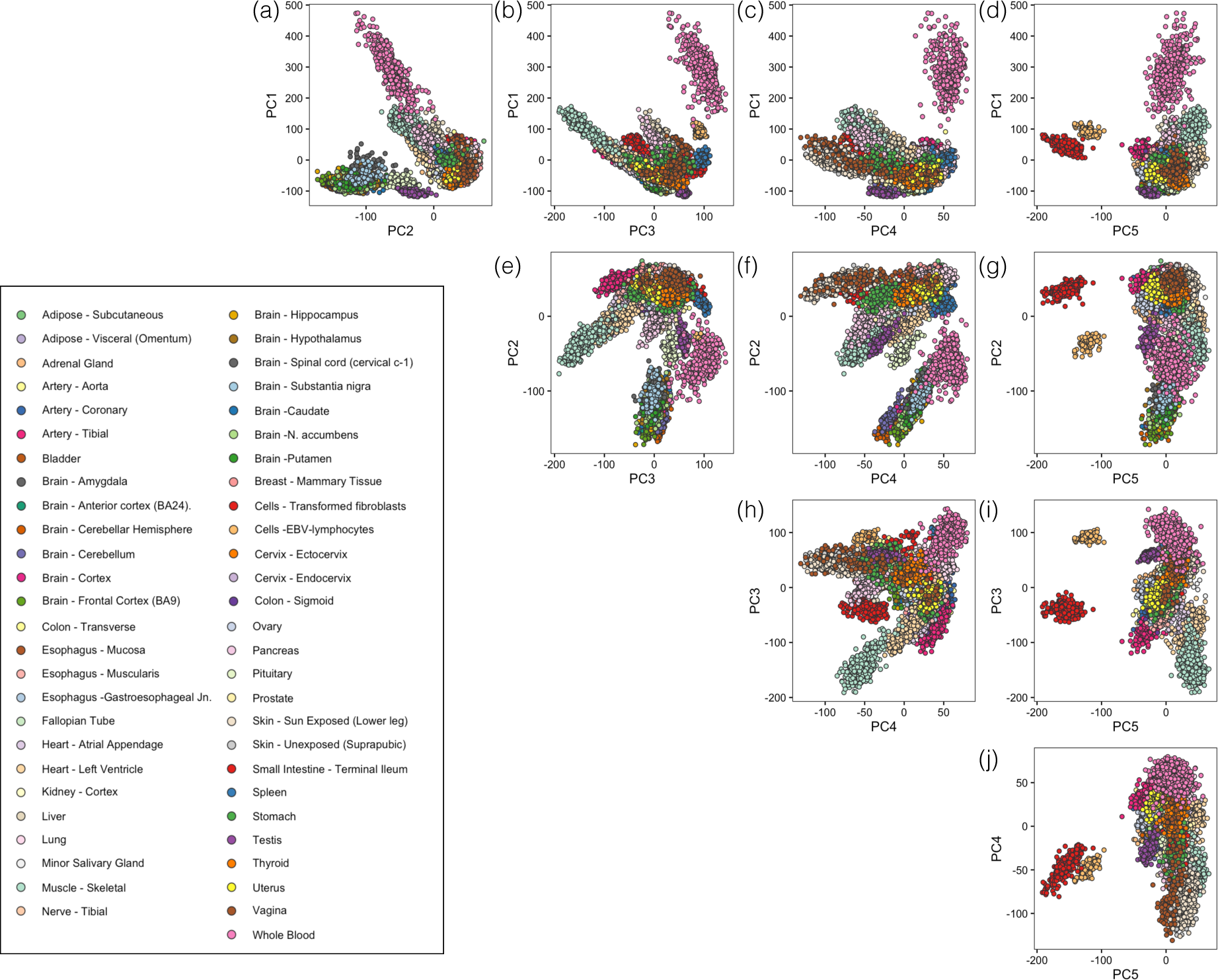
Top five principal components (PC) for GTEx V6 tissue samples. Scatter plot representation of the top five PCs of the GTEx tissue samples. Data was transformed to log2 counts per million (CPM).

**S3 Fig.**
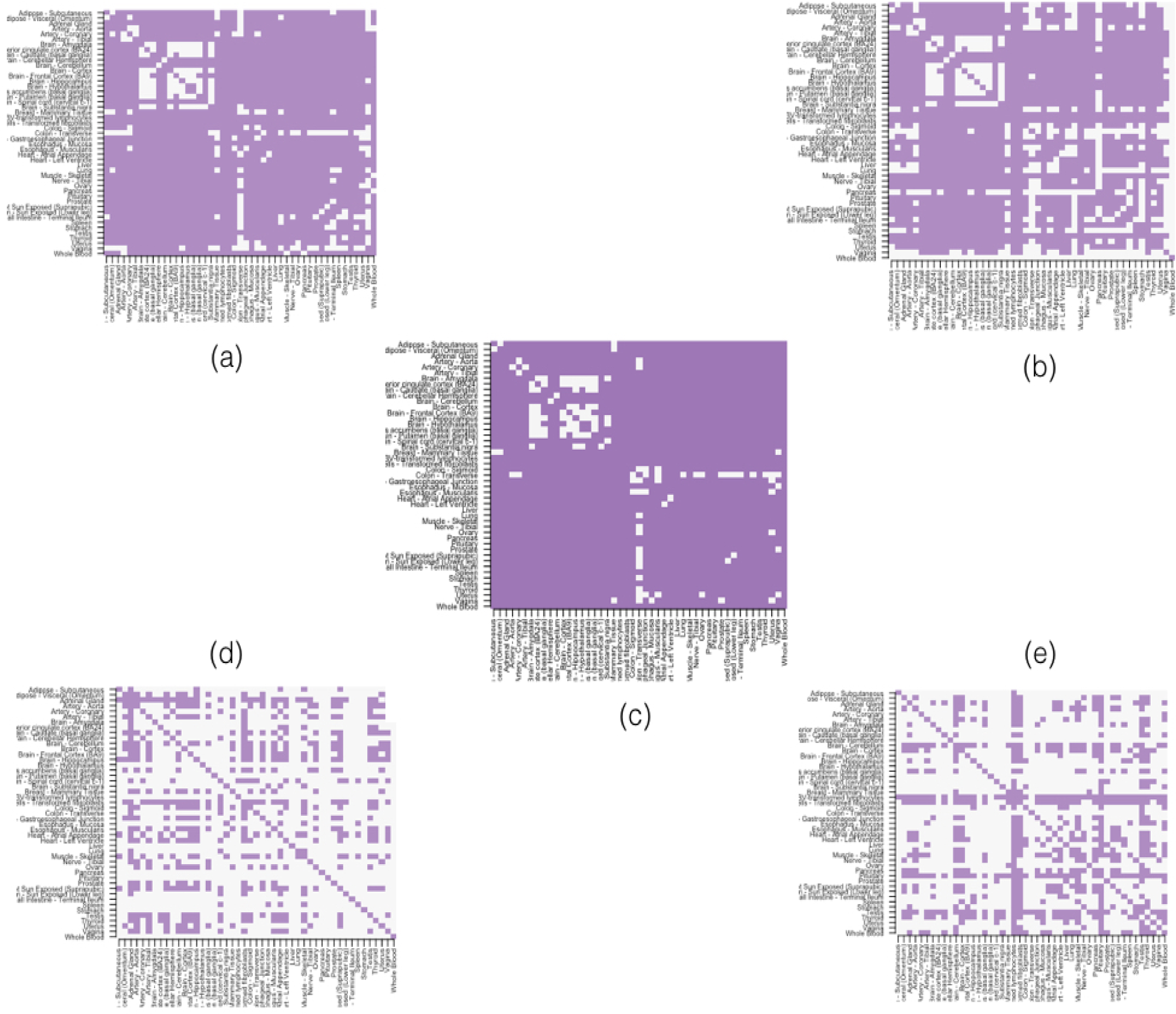
Comparison between GoM model and hierarchical clustering under different scenarios of data transformation. We used GTEx V6 data for model performance comparisons. Specifically, for every pair of the 53 tissues, we assessed the ability of the methods to separate samples according to their tissue of origin. The subplots of heatmaps show the results of evaluation under different scenarios. Filled squares in the heatmap indicate successful separation of the samples in corresponding tissue pair comparison. (a) Hierarchical clustering on log2 counts per million (CPM) transformed data using Euclidean distance. (b) Hierarchical clustering on the standardized log2-CPM transformed data (transformed values for each gene was mean and scale transformed) using the Euclidean distance. (c) GoM model of *K* = 2 applied to counts.(d) Hierarchical clustering on counts data with the assumption that, for each gene the sample read count *c_ng_* has a variance *c*¯*_g_ +* 1 that is constant across samples. And, the the gene-specific variance *c*¯*_g_ +* 1 was used to scale the distance matrix for clustering.(e) Hierarchical clustering applied to adjusted count data from c(c). Each gene expression is further normalized over (c) to have mean expression value of 0 and variance of 1. Taken together, these results suggest that regardless of the different data transformation scenarios, the GoM model with *K* = 2 is able to separate samples of different tissue of origin, better than hierarchical cluster methods.

**S4 Fig.**
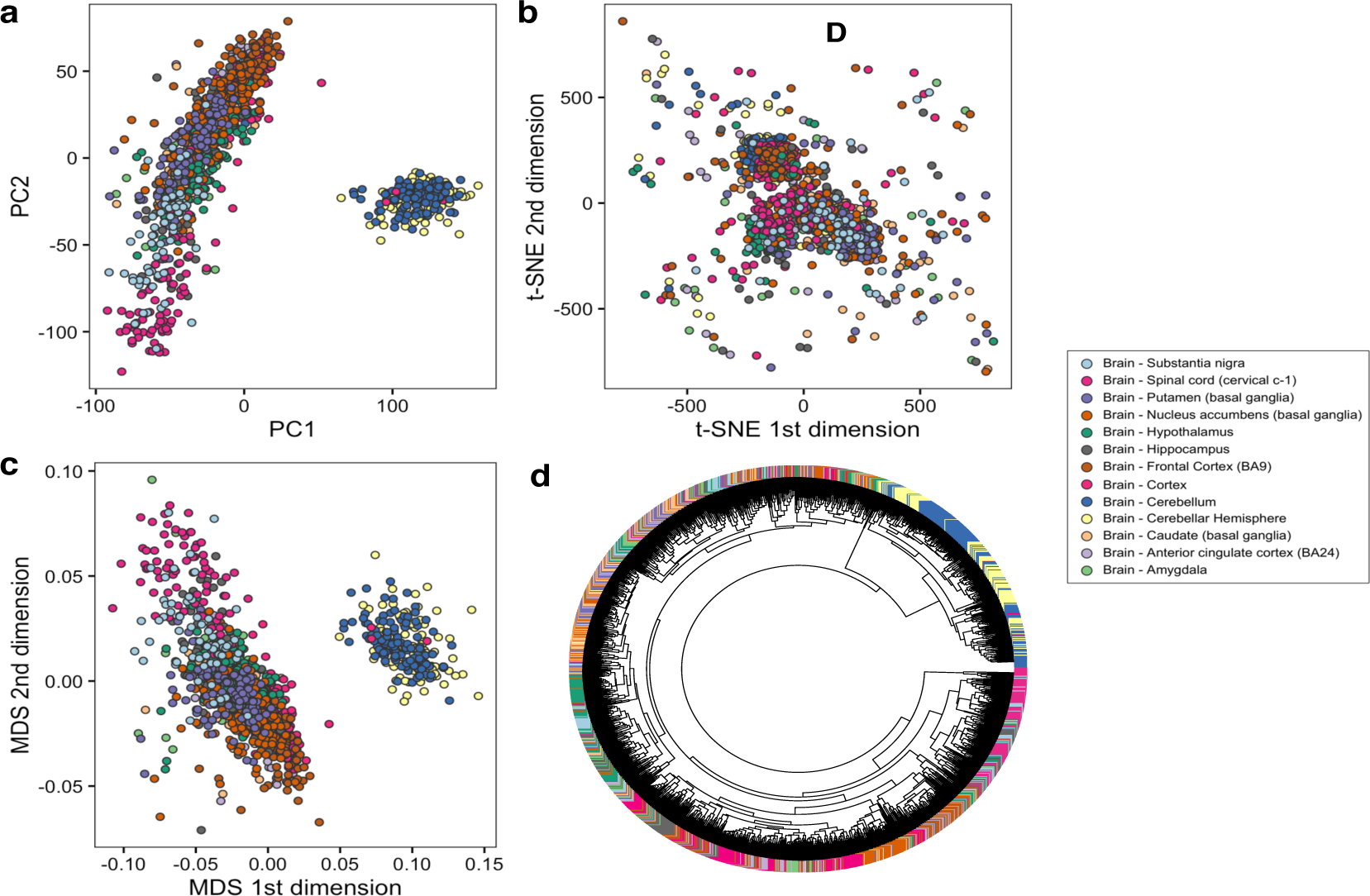
GTEx brain tissue samples visualization using (a) principle component analysis,(b) t-SNE, and (c) Multidimensional scaling and (d) dendrogram for hierarchical clustering. The colors represent the 13 different brain tissue types. In (a) and (b), the majority of the tissue samples are distinct from Cerebellum tissue samples (the cluster of samples located on the right side of the plot). While, in (c), most tissue samples are located at the enter of the plot and are similar to each other in the t-SNE dimensions. In (d), samples from Brain Cerebellar, Cerebellar Hemisphere seem to cluster together and separate from samples from other brain regions. But, because of the large number of samples, patterns of variation between tissue samples are difficult to detect.

**S5 Fig.**
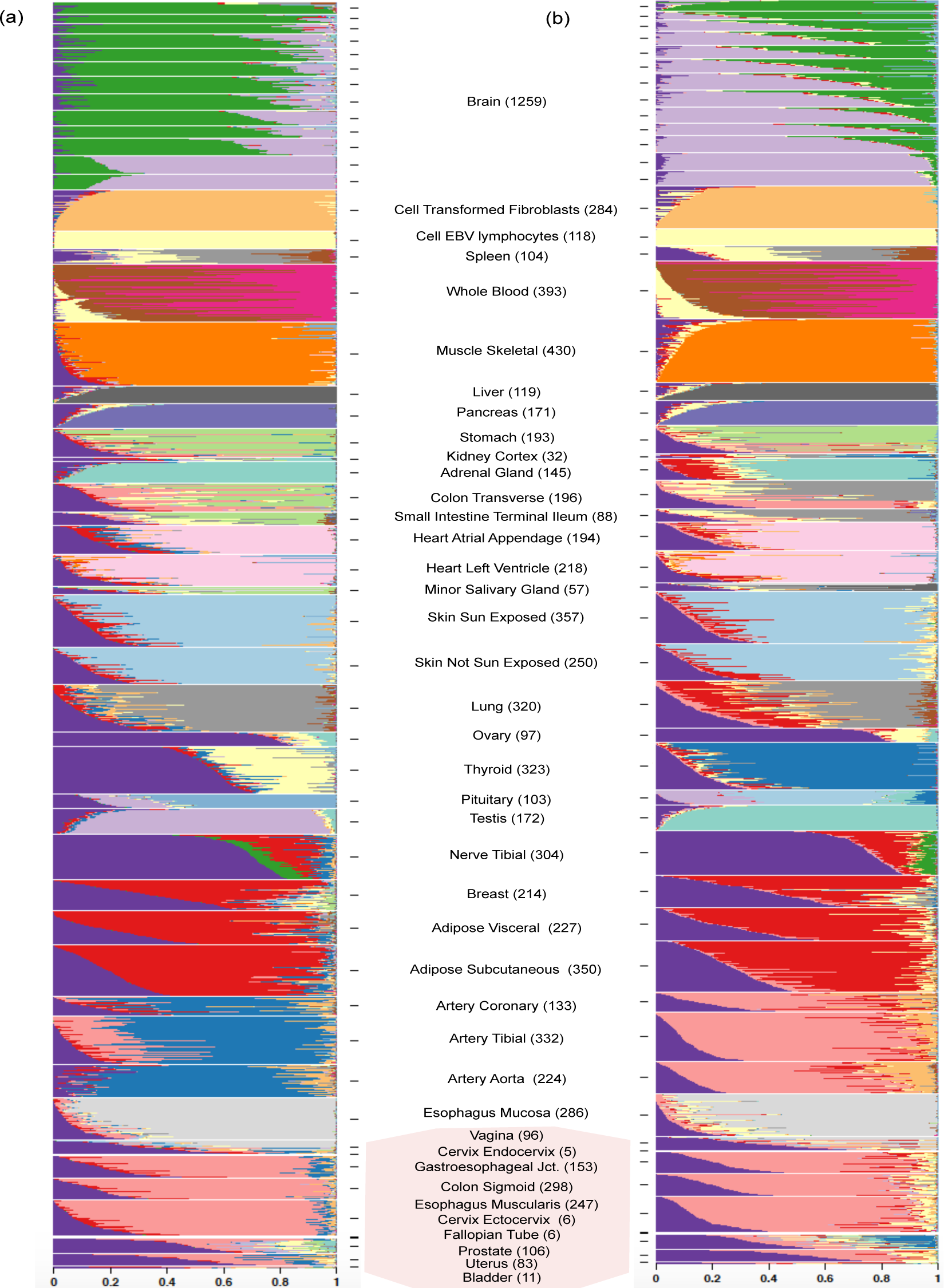
Structure plot of GTEx V6 tissue samples for *K* = 20 in two runs under different thinning parameter settings. (a) *p_thin_ =* 0.01 and (b) *p_thin_ =* 0.0001. The structural patterns in these two plots closely resemble the structural patterns in Fig 1(a), though there are a few differences from the unthinned version.

**S6 Fig.**
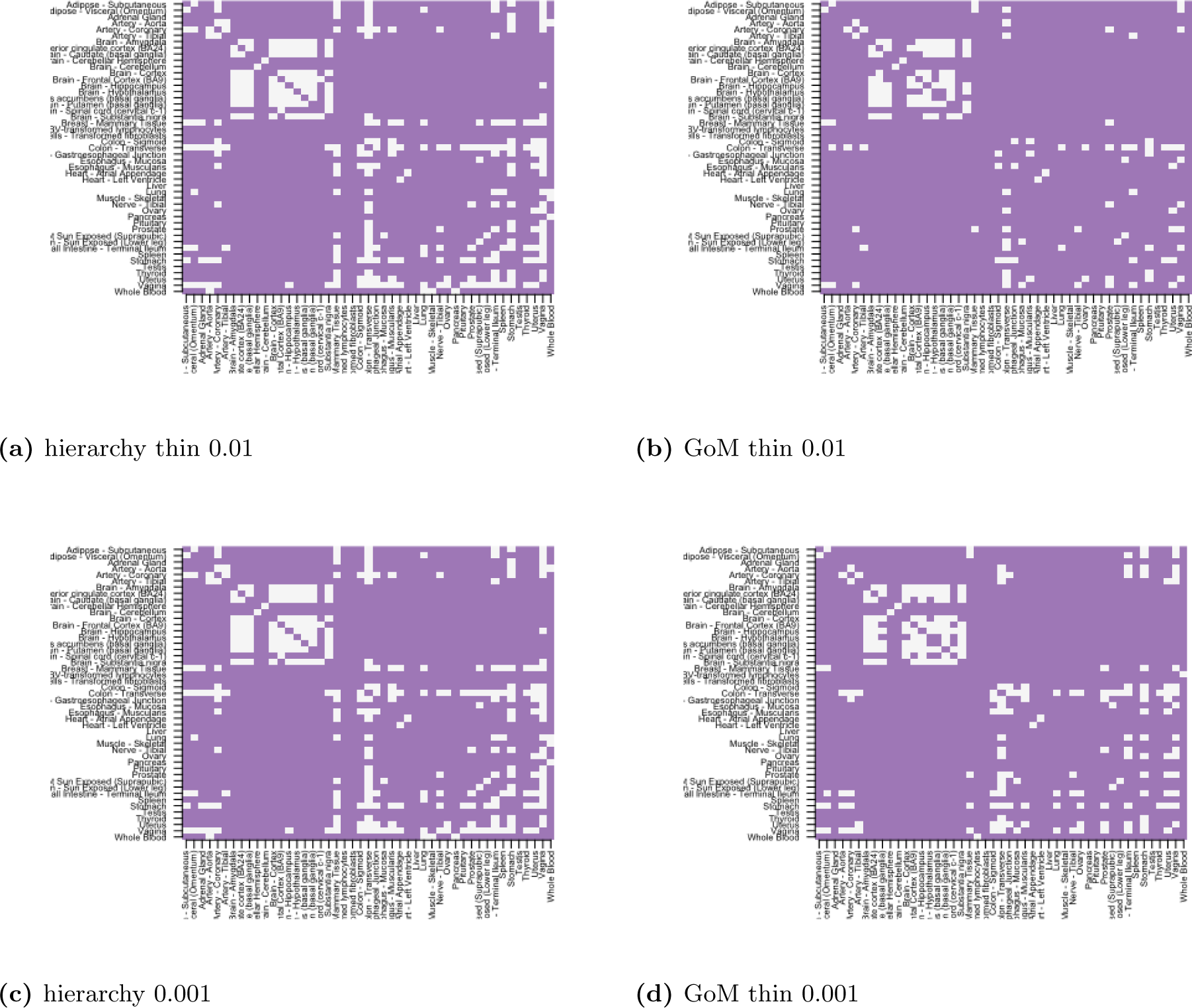
A comparison of “accuracy” of hierarchical clustering vs. GoM on thinned GTEx data, with thinning parameters of *p_thin_ =* 0.01 and *p_thin_ =* 0.001. For each pair of tissue samples from the GTEx V6 data we assessed whether or not each clustering method (with *K* = 2 clusters) separated the samples according to their tissue of origin, with successful separation indicated by a filled square. Thinning deteriorates accuracy compared with the unthinned data (Fig 2), but even then the model-based method remains more successful than the hierarchical clustering in separating the samples by tissue or origin.

**S7 Fig.**
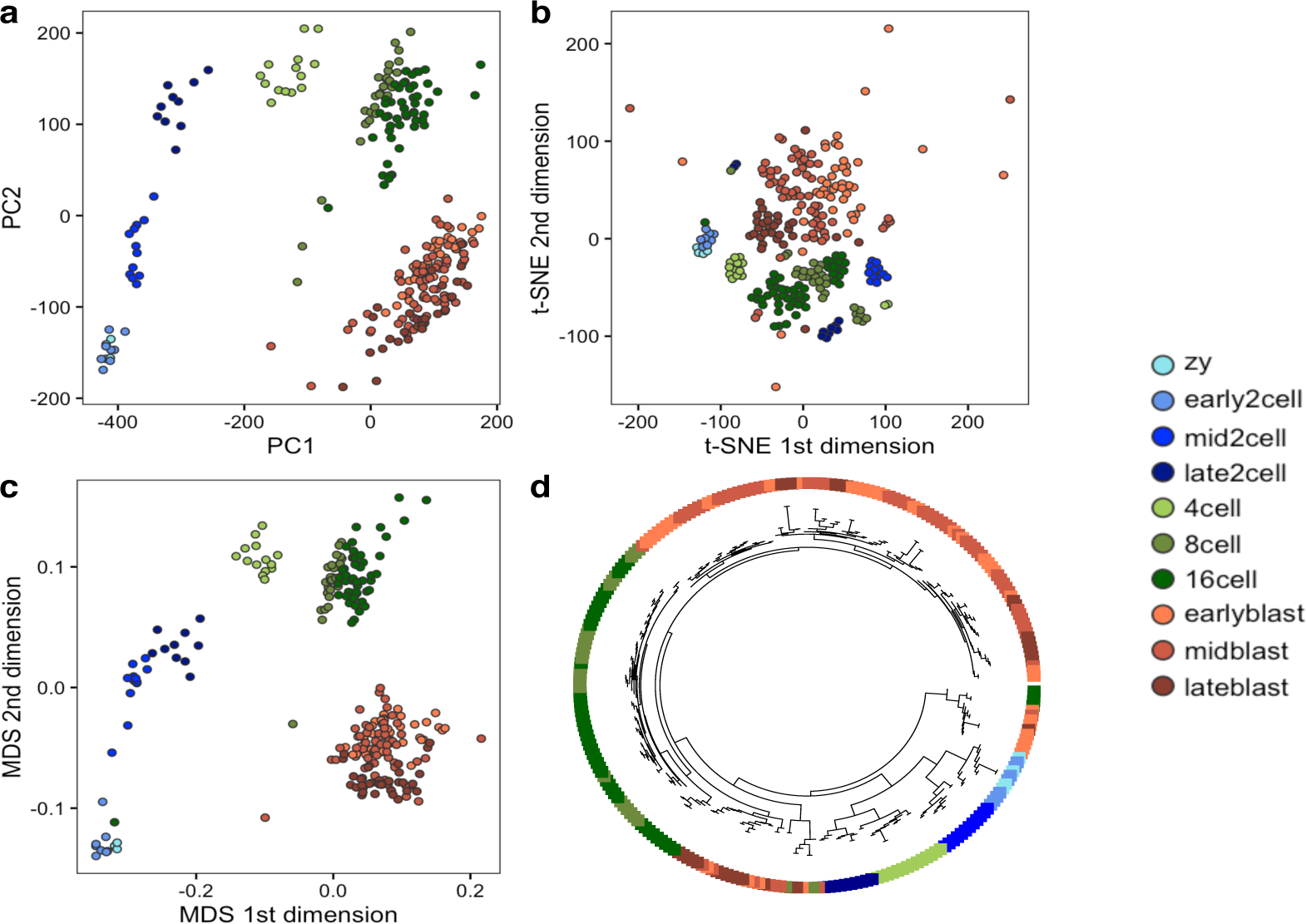
Visualizing mouse pre-implantation embryos data from Deng et al (2014) using (a) Principle Component Analysis, (b) t-SNE, (c) Multidimensional Scaling (MDS) and (d) circular dendrogram for hierarchical clustering. The colors represent different developmental stages. PCA, MDS seem to be effective in capturing the developmental trajectory, but t-SNE fails to do so. Hierarchical clustering fails to separate out the blastocyst cells from the cells in early stages of development completely.

**S8 Fig.**
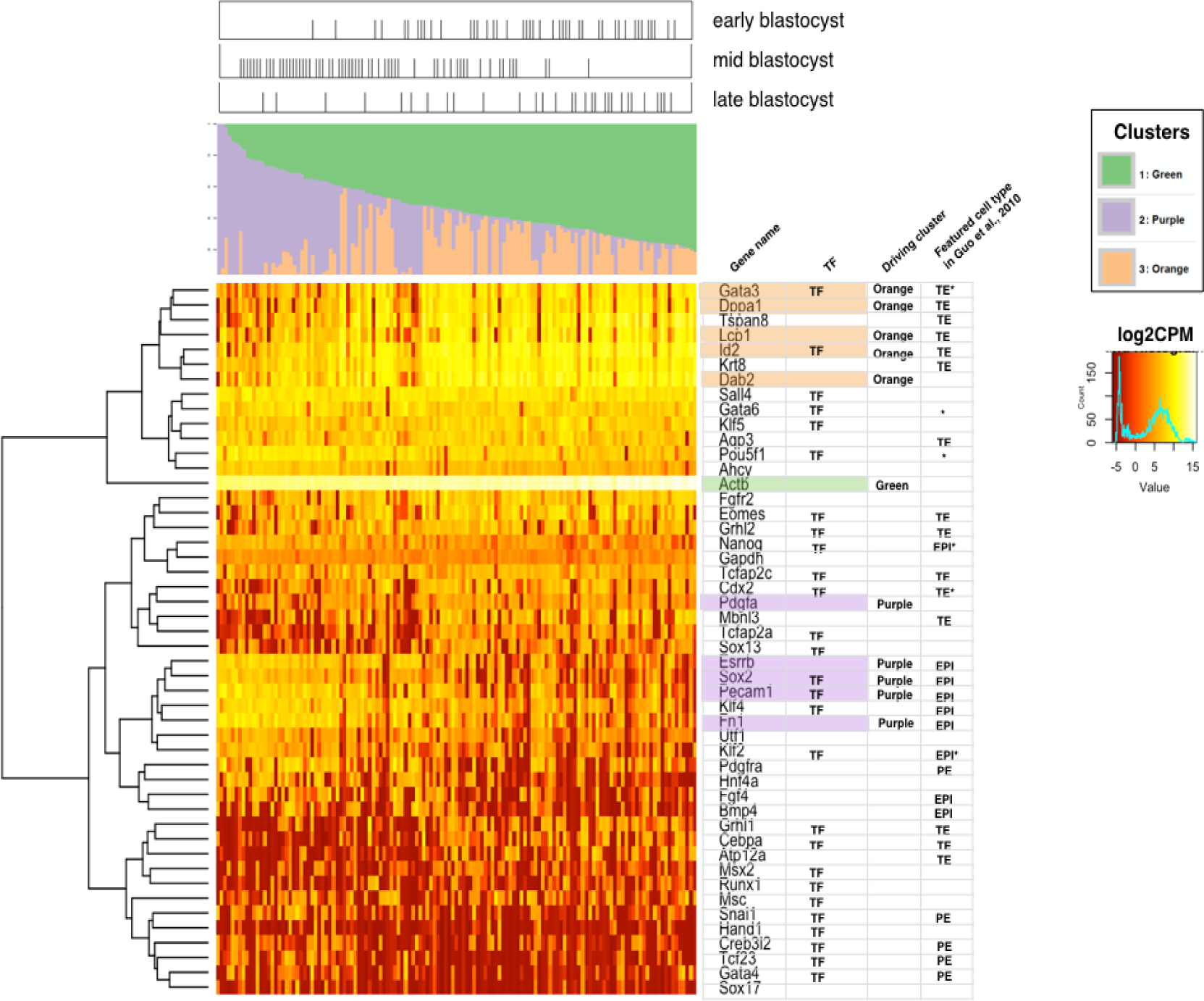
Additional GoM analysis of Deng et al (2014) data including blastocyst samples and 48 blastocyst marker genes. We considered 48 blastocyst marker genes (as chosen by Guo et al., 2010) and fitted GoM model with *K* = 3 to 133 blastocyst samples. In the Structure plot, blastocyst samples are arranged in order of estimated membership proportion in the Green cluster. The panel located above the Structure plot shows the corresponding pre-implantation stage from which blastocyst samples were collected. The heatmap located below the Structure plot represents expression levels of the 48 blastocyst marker genes (log2 CPM), and the corresponding dendrogram shows results of hierarchical clustering (complete linkage). The table on the right of the expression heatmap displays gene information, showing, from left to right, 1) whether or not the gene is a transcription factor, 2) the driving GoM cluster if the gene was among the top five driving genes, and 3) the featured cell type (TE: trophecoderm, EPI: epiblast, PE: primitive endoderm) that was found in Guo et al., 2010.

**S9 Fig.**
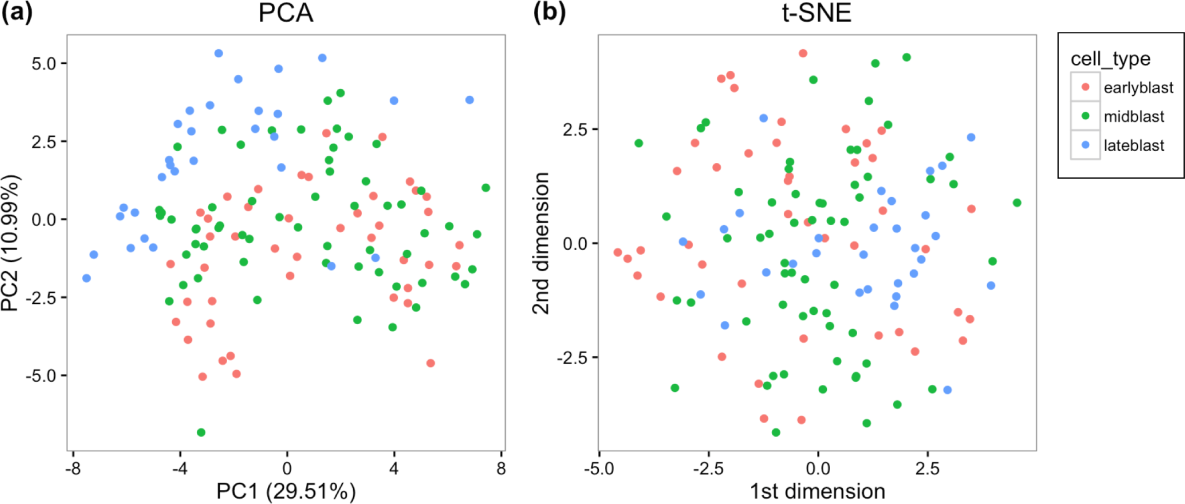
Visualization of PCA and t-SNE results of mouse pre-implantation embryos data from Deng et al (2014) using 48 blastocyst marker genes

**S10 Fig.**
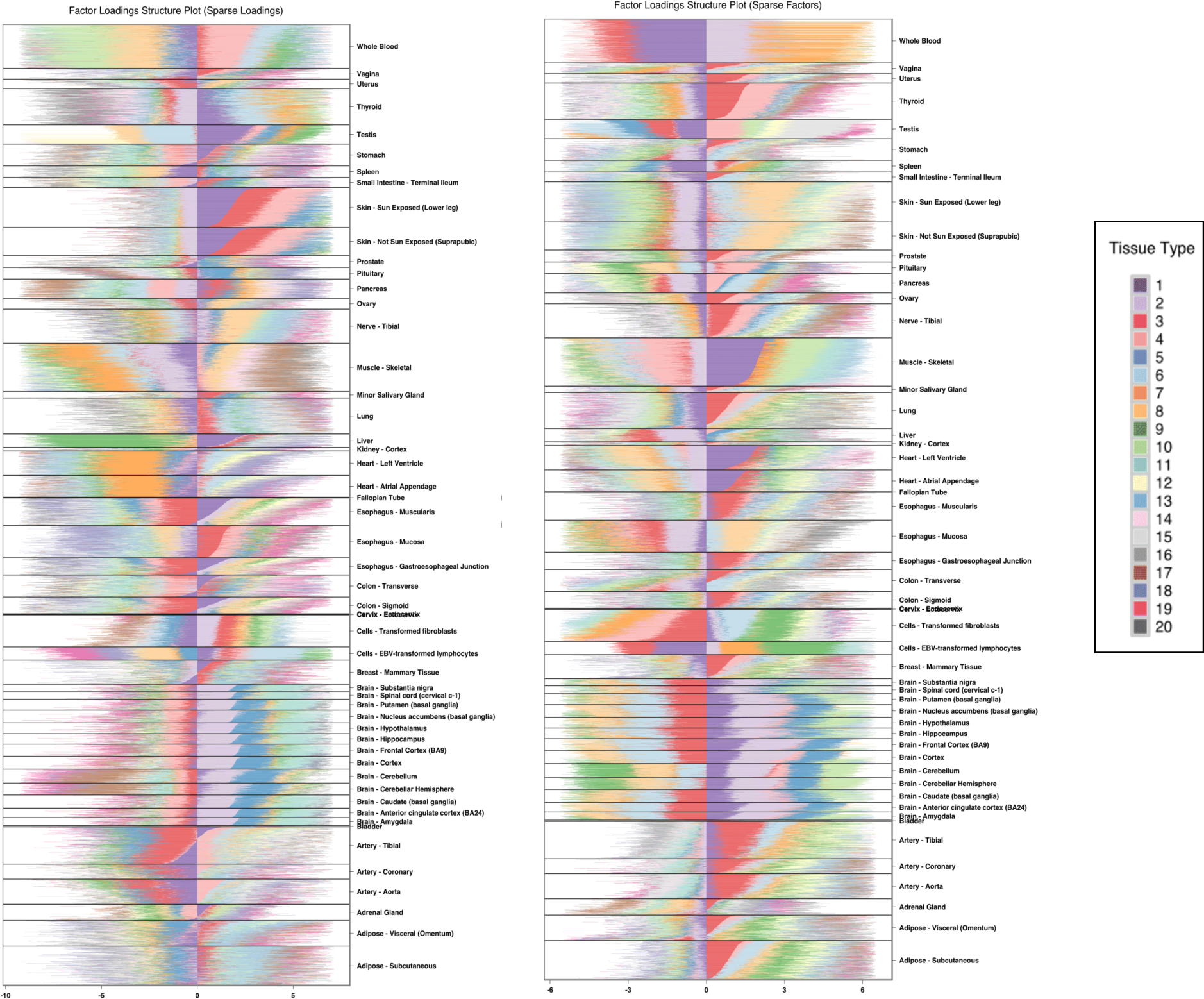
Sparse Factor Analysis loadings visualization of GTEx V6 tissue samples. The colors represent the 20 different factors. The factor loadings are presented in a stacked bar for each sample. We performed SFA under the scenarios of (left) when the loadings are sparse and (right) when the factors are sparse.

**S11 Fig.**
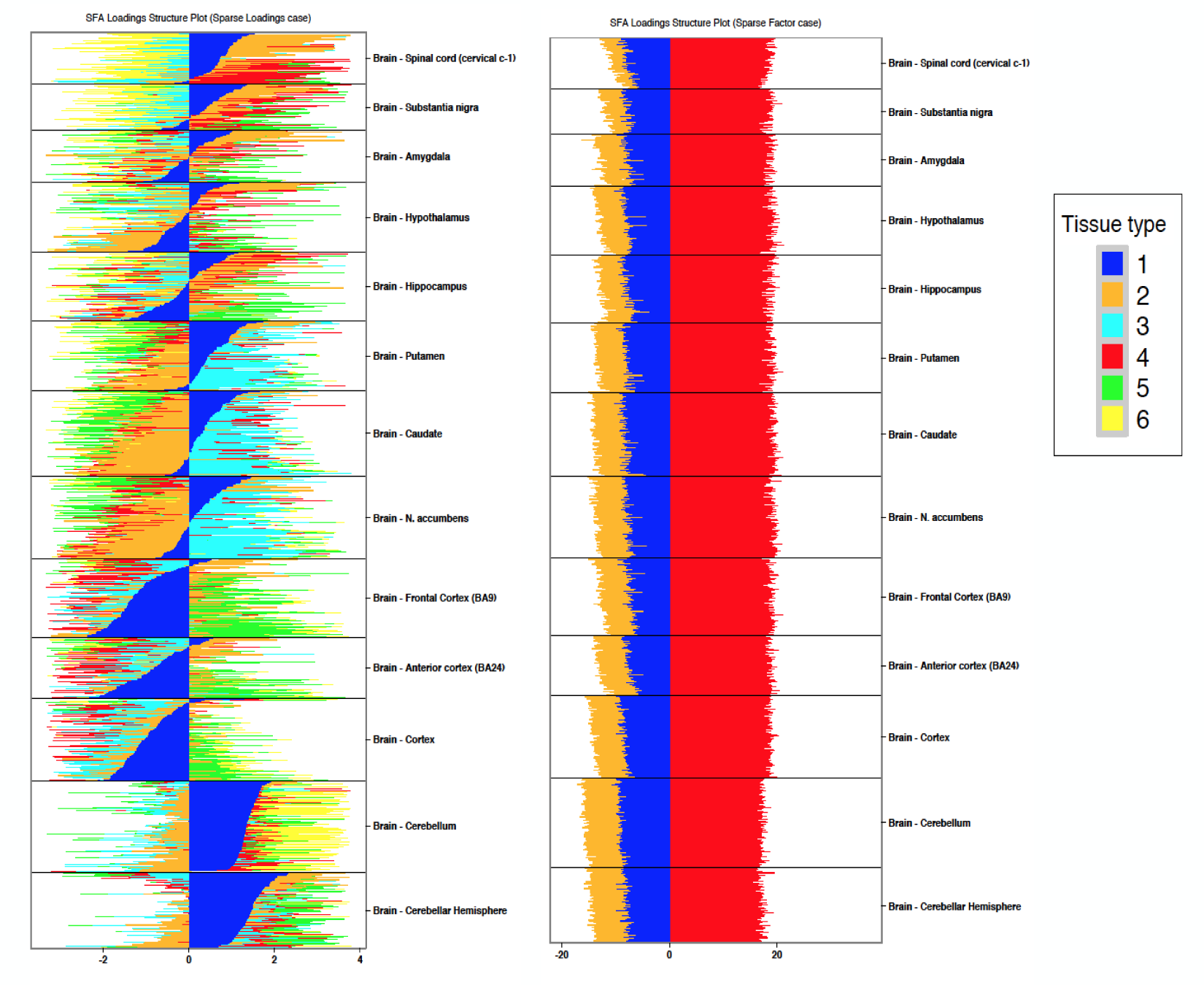
Sparse Factor Analysis loadings visualization of GTEx brain tissue samples. The colors represent the 6 different factors. The factor loadings are presented in a stacked bar for each sample. We performed SFA under the scenarios of (left) when the loadings are sparse and (right) when the factors are sparse.

**S12 Fig.**
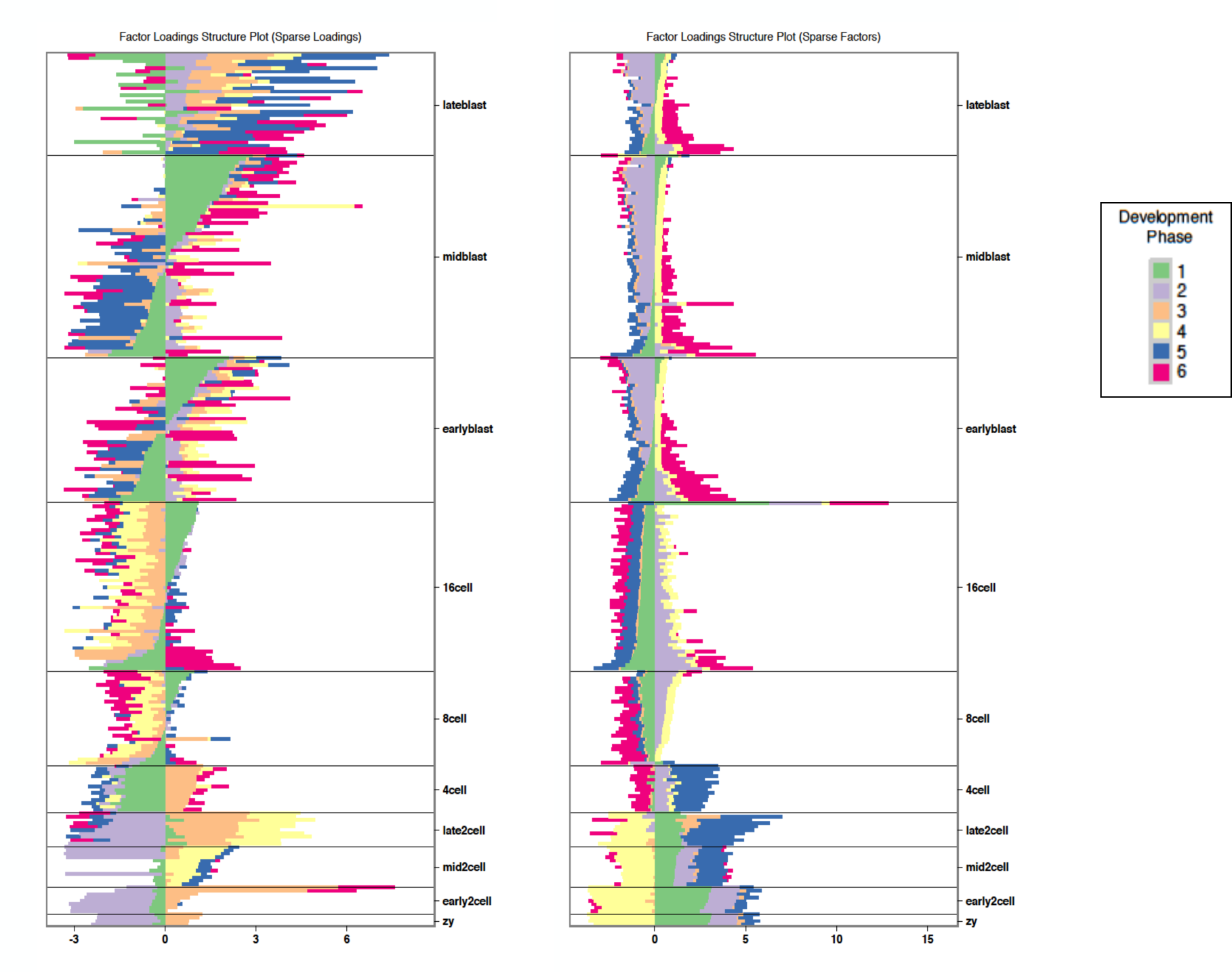
Sparse Factor Analysis loadings visualization of mouse pre-implantation embryos from Deng et al., (2014). The colors represent the 6 different factors. The factor loadings are presented in a stacked bar for each sample. We performed SFA under the scenarios of (left) when the loadings are sparse and (right) when the factors are sparse.

## Supplementary tables

**S1 Table.**
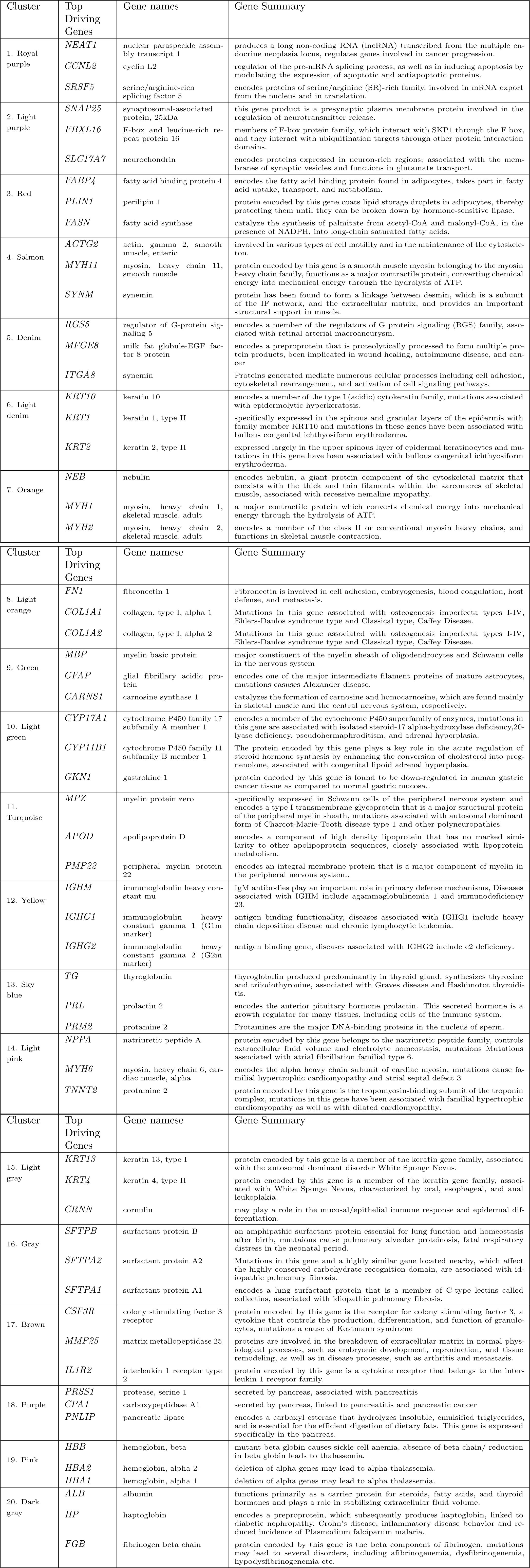
Cluster Annotations of GTEx V6 data with top driving gene summaries.

**S2 Table.** Cluster Annotations of GTEx V6 Brain data with top driving gene summaries.

| Cluster | Top Driving Genes | Gene names | Gene Summary |
| --- | --- | --- | --- |
| 1, Royal blue | *CLU* | clusterin | protein encoded by this gene is a secreted chaperone that can under some stress conditions also be found in the cell cytosol, also involved in cell death, tumor progression, and neurodegenerative disorders. |
|  | *OXT* | oxytocin/neurophysin I pre-propeptide | encodes a precursor protein that is processed to produce oxytocin and neurophysin I, involved in contraction of smooth muscle during parturition and lactation, cognition, tolerance, adaptation and complex sexual and maternal behaviour. |
|  | *GLUL* | glutamate-ammonia ligase | catalyzes the synthesis of glutamine from glutamate and ammonia in an ATP-dependent reaction, associated with congenital glutamine deficiency, and overexpression of this gene was observed in some primary liver cancer samples. |
| 2, Turquoise | *ENC1* | ectodermal-neural cortex 1 | plays a role in the oxidative stress response as a regulator of the transcription factor Nrf2, may play role in malignant transformation. |
|  | *NCALD* | neurocalcin delta | encodes a member of the neuronal calcium sensor (NCS), a regulator of G protein-coupled receptor signal transduction. |
|  | *YWHAH* | tyrosine 3-monooxygenase/tryptophan 5-monooxygenase activation protein eta | mediate signal transduction by binding to phosphoserine-containing proteins, associated with early-onset schizophrenia and psychotic bipolar disorder. |
| 3, Lime green | *PKD1* | polycystin 1, transient receptor potential channel interacting | functions as a regulator of calcium permeable cation channels and intracellular calcium homeostasis. It is also involved in cell-cell/matrix interactions and may modulate G-protein-coupled signal-transduction pathways. |
|  | *CBLN3* | cerebellin 3 precursor | contain a cerebellin motif and C-terminal C1q signature domain that mediates trimeric assembly of atypical collagen complexes |
|  | *CHGB* | chromogranin B | encodes a tyrosine-sulfated secretory protein abundant in peptidergic endocrine cells and neurons. This protein may serve as a precursor for regulatory peptides. |
| 4, Red | *PPP1R1B* | protein phosphatase 1 regulatory inhibitor sub- unit 1B | encodes a bifunctional signal transduction molecule, may serve as a therapeutic target for neurologic and psychiatric disorders. |
|  | *RGS14* | regulator of G-protein signaling 14 | attenuates the signaling activity of G-proteins, increases the rate of conversion of the GTP to GDP. |
|  | *NCDN* | neurochondrin | encodes a leucine-rich cytoplasmic protein, essential for spatial learning processes. |
| 5, Yellow orange | *MBP* | myelin basic protein | protein encoded is a major constituent of the myelin sheath of oligodendrocytes and Schwann cells in the nervous system. |
|  | *GFAP* | glial fibrillary acidic protein | encodes major intermediate filament proteins of mature astrocytes, a marker to distinguish astrocytes during development, mutations in this gene cause Alexander disease, a rare disorder of astrocytes in central nervous system. |
|  | *TF* | transferrin | transport iron from the intestine, reticuloendothelial system, and liver parenchymal cells to all proliferating cells in the body, involved in the removal of certain organic matter and allergens from serum. |
| 6, Yellow | *IQGAP1* | IQ motif containing GTPase activating protein 1 | interacts with components of the cytoskeleton, with cell adhesion molecules, and with several signaling molecules to regulate cell morphology and motility. |
|  | *A2M* | alpha-2-macroglobulin | inhibits many proteases, including trypsin, thrombin and collagenase. A2M is implicated in Alzheimer disease (AD) due to its ability to mediate the clearance and degradation of A-beta, the major component of beta-amyloid deposits. |
|  | *C3* | complement component 3 | plays a central role in the activation of complement system, associated with atypical hemolytic uremic syndrome and age-related macular degeneration in human patients. |

**S3 Table.** Cluster Annotations of Deng data with top driving genes.

|  | GO ID | GO Term | Top Driving Genes |
| --- | --- | --- | --- |
| 1 | GO:0007276 | gamete generation | BCL2L10; GDF9; NOBOX; PABPC1L; RGS2; CREB3L4; RNF114; BMP15; PTTG1; TDRD12; WEE2; SPIN1; DAZL |
| 2 | GO:0007292 | female gamete generation | GDF9; BCL2L10; PABPC1L; BMP15; WEE2; DAZL; NOBOX |
| 3 | GO:0048609 | multicellular organismal reproductive process | GDF9; NOBOX; PABPC1L; BCL2L10; BMP15; CREB3L4; TGFB2; RNF114; RGS2; PTTG1; TDRD12; WEE2; SPIN1; DAZL |
| 4 | GO:0032504 | multicellular organism reproduction | GDF9; NOBOX; PABPC1L; BCL2L10; BMP15; CREB3L4; TGFB2; RNF114; RGS2; PTTG1; TDRD12; WEE2; SPIN1; DAZL |
| 5 | GO:0019953 | sexual reproduction | BCL2L10; GDF9; NOBOX; PABPC1L; RGS2; CREB3L4; RNF114; BMP15; PTTG1; TDRD12; WEE2; SPIN1; DAZL |
| 6 | GO:0044702 | single organism reproductive process | GDF9; NOBOX; PABPC1L; BCL2L10; BMP15; CREB3L4; TGFB2; CASP8; RNF114; RGS2; PTTG1; TDRD12; WEE2; SPIN1; DAZL |
| 7 | GO:0048477 | oogenesis | WEE2; GDF9; NOBOX; PABPC1L; DAZL |
| 8 | GO:0044703 | multi-organism reproductive process | BCL2L10; GDF9; NOBOX; PABPC1L; RGS2; CREB3L4; RNF114; BMP15; PTTG1; TDRD12; WEE2; SPIN1; DAZL |
| 9 | GO:0048599 | oocyte development | WEE2; GDF9; PABPC1L; DAZL |
| 10 | GO:0009994 | oocyte differentiation | WEE2; GDF9; PABPC1L; DAZL |
| 11 | GO:0051321 | meiotic cell cycle | H1FOO; WEE2; TDRD12; SPIN1; PTTG1; DAZL |
| 12 | GO:0001556 | oocyte maturation | WEE2; PABPC1L; DAZL |
| 13 | GO:0006306 | DNA methylation | TDRD12; H1FOO; TET3; ZFP57 |
| 14 | GO:0051302 | regulation of cell division | TGFB2; PTTG1; TXNIP; WEE2; CHEK1; DAZL |
| 15 | GO:0060255 | regulation of macromolecule metabolic process | TGFB2; NOBOX; BPGM; UBE2D3; NFYA; CASP8; BMP15; TXNIP; TDRD12; GDF9; BCL2L10 |

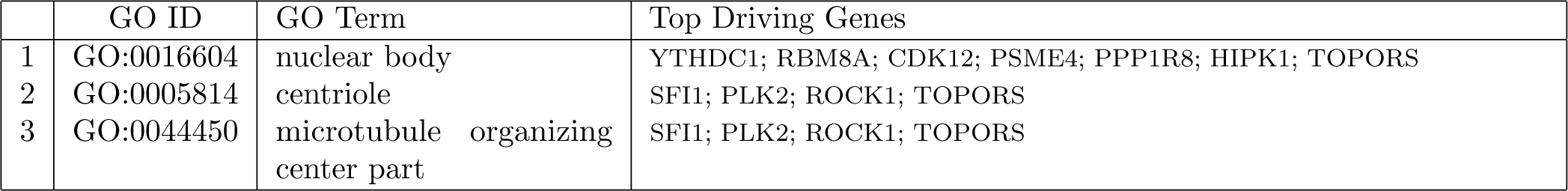
Deng et al (2014) Cluster 2 (magenta) top GO annotations.

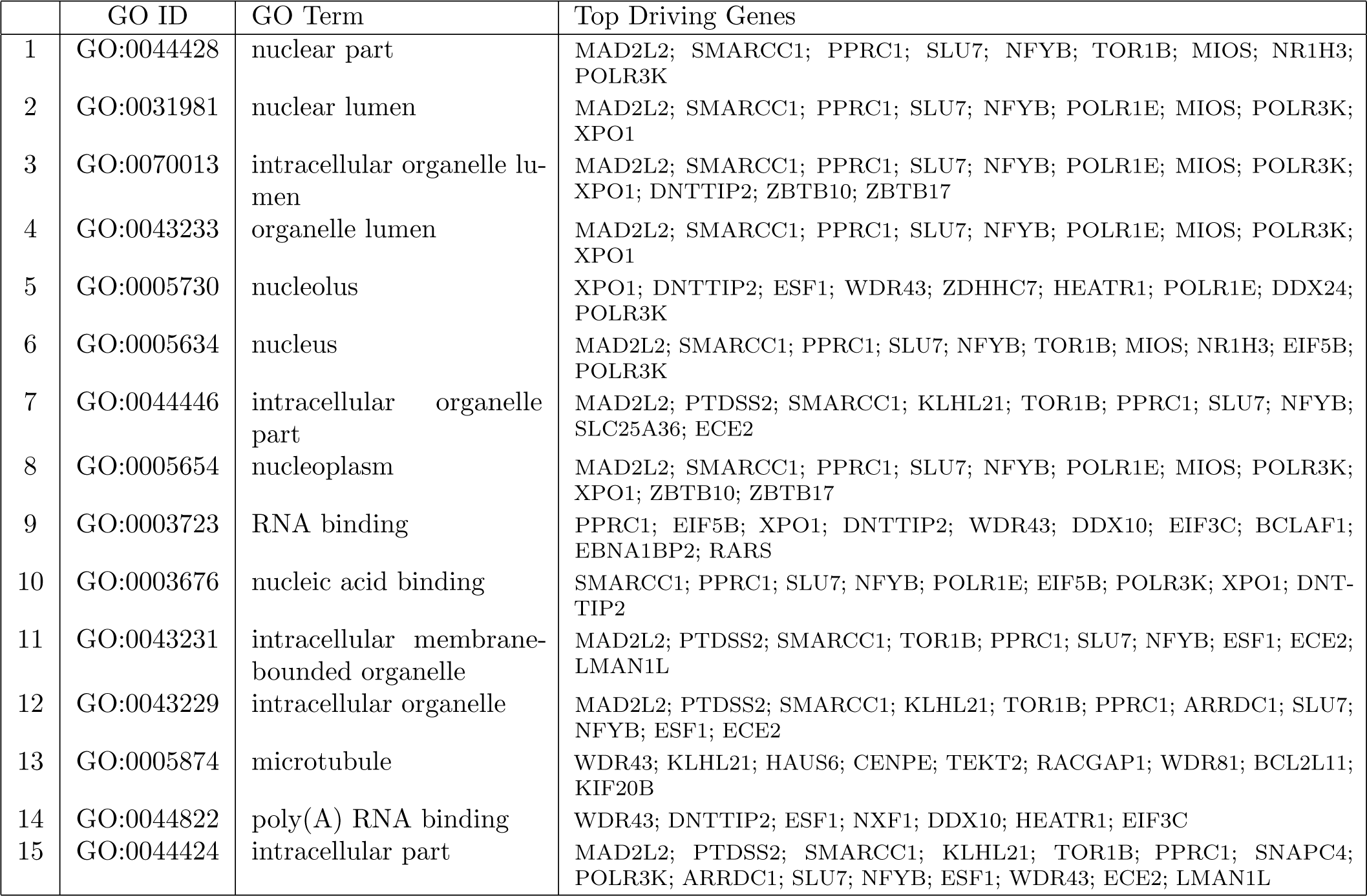
Deng et al (2014) Cluster 3 (yellow) top GO annotations.

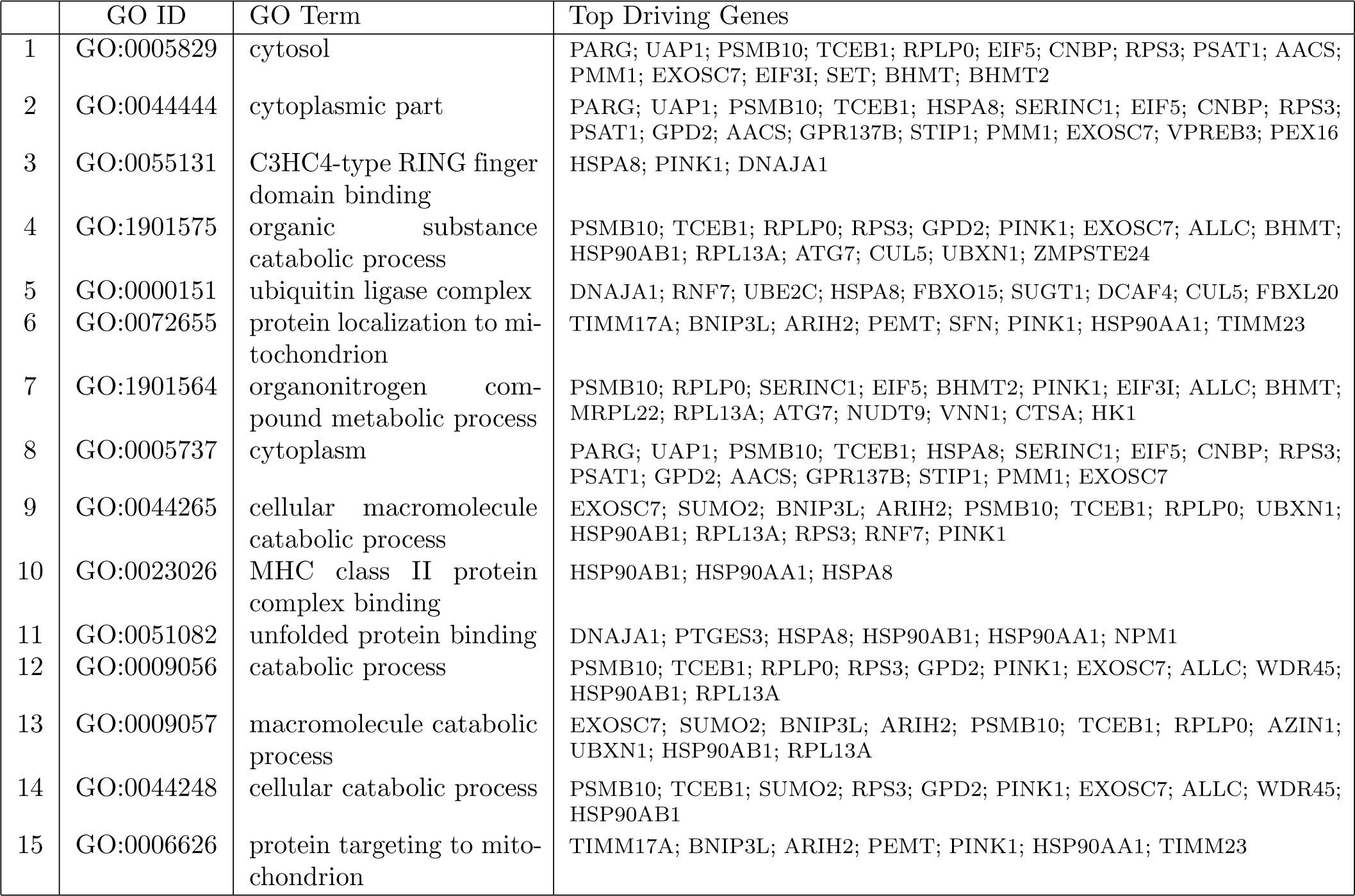
Deng et al (2014) Cluster 4 (green) top GO annotations.

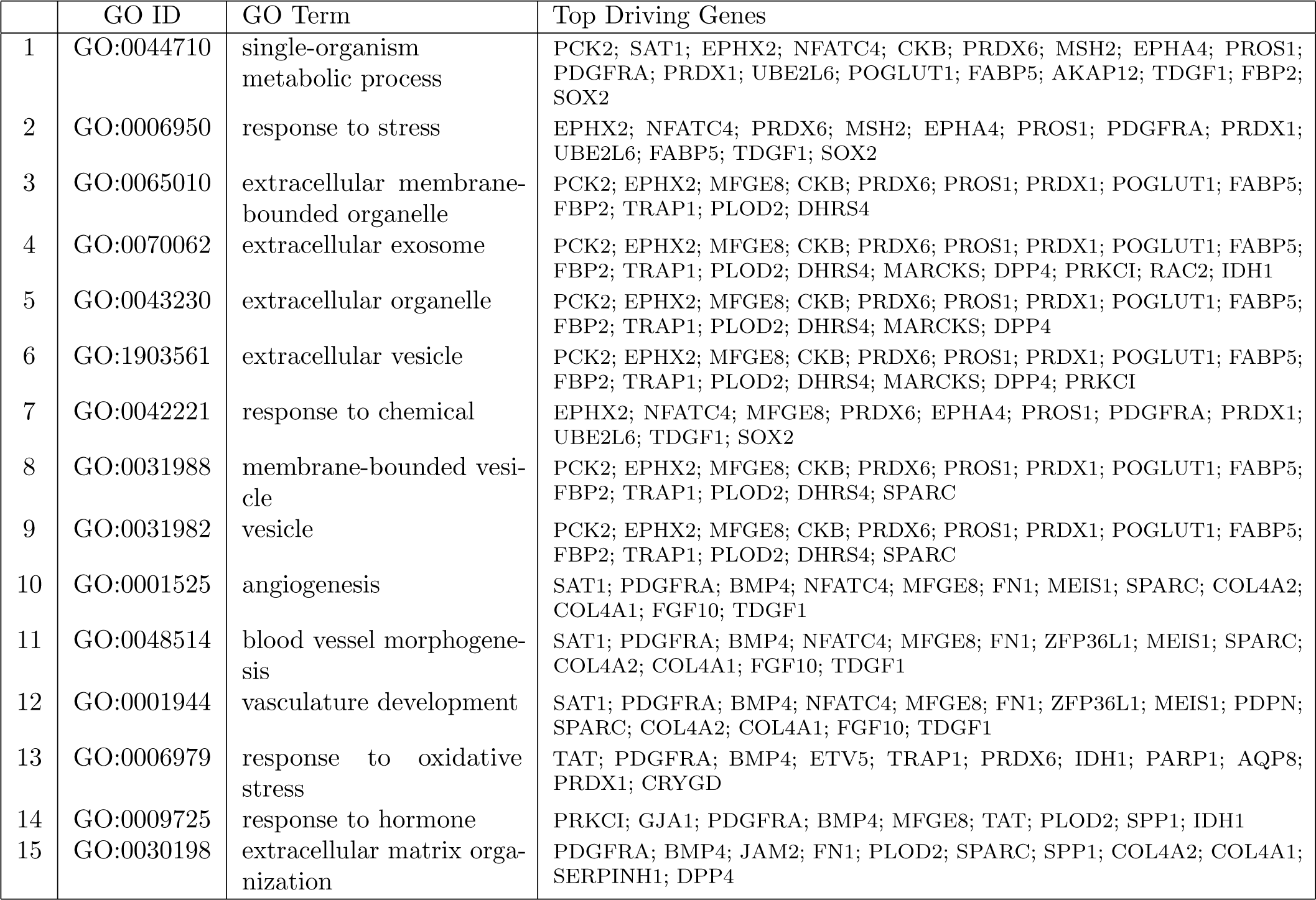
Deng et al (2014) Cluster 5 (purple) top GO annotations.

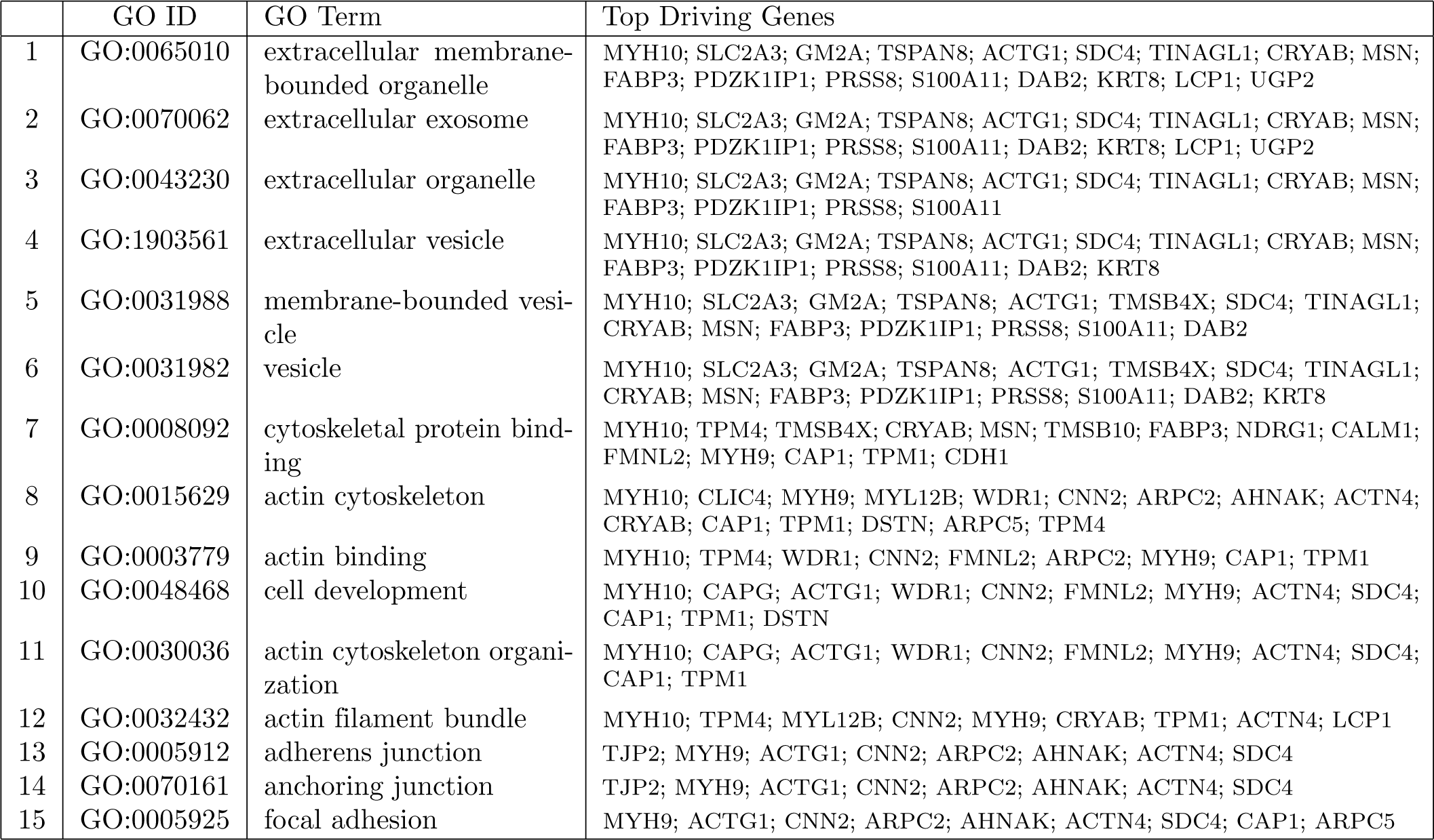
Deng et al (2014) Cluster 6 (orange) top GO annotations.

**S4 Table.** Cluster Annotation of Deng data analysis using 48 genes with top driving gene summaries.

| Cluster | Top 5 Driving Genes | Top significant GO terms (function)[q-value] |
| --- | --- | --- |
| Green | *Actb* | GO:0048568 (embryonic organ development)[9e-08], GO:0048468 (cell development)[4e-07], GO:0001890 (placenta development)[1e-06], GO:0051094 (positive regulation of developmental process)[1e-06], GO:0030097 (hemopoiesis)[1e-05] |
| Purple | *Pecam1, Esrrb, Fn1, Pdgfra, Sox2* | GO:0048864 (stem cell development)[4e-12], GO:0048863 (stem cell differentiation)[2e-11], GO:0009893 (positive regulation of metabolic process)[7e-10], GO:0009653 (anatomical structure morphogenesis)[4e-08] |
| Orange | *Dppa1, Gata3, Id2, Dab2, Lcp1* | GO:0061061 (muscle structure development)[2e-13], GO:0060537 (muscle tissue development)[2e-12], GO:0048514 (blood vessel morphogenesis)[8e-12], GO:0007275 (multicellular organismal development) |

